# DeepAden: An explainable machine learning method for predicting the substrate specificity of nonribosomal peptide synthetases

**DOI:** 10.1101/2025.05.21.655435

**Authors:** Jiaquan Huang, Liangjun Ge, Yaxin Wu, Song Meng, Yi Tian, Jun Wu, Qiandi Gao, Pan Li, Heqian Zhang, Zhiwei Qin

## Abstract

Microbial non-ribosomal peptides (NRPs) exhibit remarkable structural diversity and serve as valuable sources of lead compounds for clinical drug development. The biosynthesis of NRPs relies on non-ribosomal peptide synthetases (NRPSs), in which adenylation (A) domains play a pivotal role in defining the core structure by selectively recognizing and activating amino acid substrates. Accurately predicting the substrate specificities of A-domains is thus essential for understanding the core structural and biosynthetic logic of NRPs. Here, we present DeepAden, a two-stage deep learning framework. In the first stage, a graph attention network (GAT)-based model localizes 27-residue binding pockets within 6 Å of bound substrates and convert these into pocket representations. In the second stage, pocket representations are then encoded alongside substrate information using pretrained language models, and aligned using contrastive learning. In addition, we introduce a SHapley Additive exPlanations (SHAP)-guided data augmentation strategy to mitigate class imbalance and improve robustness, particularly for nonproteinogenic substrates. DeepAden achieves competitive performance compared with state-of-the-art tools on a benchmark dataset, and enabled the identification of two *Streptomyces* NRPS gene clusters through accurate A-domain substrates specificity predictions. DeepAden offers a powerful tool for precise pocket localization and robust substrate prediction, accelerating the discovery and characterization of novel NRP natural products for future work. The DeepAden web server is available at https://deepnp.site/.

## 1. Introduction

Nonribosomal peptides (NRPs) are naturally occurring biopolymers that exhibit remarkable structural and functional diversity, acting as antibiotics, siderophores, toxins and immunosuppressants[1]. The assembly processes of their peptide backbones are mediated by highly modular megasynthetases, termed nonribosomal peptide synthetases (NRPSs), which utilize amino acids as monomers (commonly referred to as building blocks)[2]. NRPSs are minimally composed of three core domains: an adenylation (A) domain, which is responsible for substrate selection and activation; an peptidyl carrier protein (PCP) domain, which tethers activated amino acids via its phosphopantetheinyl prosthetic group; and a condensation (C) domain, which catalyses the peptide bond formation procedure to elongate the chain[3]. In principle, the sequential arrangement of A-domains within biosynthetic gene clusters (BGCs) determines the core peptide sequence. Therefore, understanding their specificities is critical for predicting peptide structures[4].

In 1997, Stachelhaus et al. reported the first crystal structure of a NRPS adenylation domain in complex with its substrate phenylalanine (GrsA-Phe, PDB ID: 1amu)[5]. Based on this structural insight, they identified 10 key amino acid residues (10-AA) within the active site that interact directly with the substrate and proposed that these specificity-conferring codes (SCCs) could be used to predict A-domain substrate specificity. Building on this concept, Rausch et al. introduced an extended A-domain binding pocket (ABP), consisting of 34 amino acid residues (34-AA) located within an 8 Å radius of the substrate[6]. The SCC and ABP frameworks have since underpinned the development of numerous A-domain substrate prediction tools, employing diverse computational approaches such as sequence alignment, hidden Markov models (HMMs)[7], support vector machines (SVMs)[8], and random forests (RF)[9]. Such tools including NRPSpredictor2[10], SANDPUMA[11], AdenPredictor[12], and PARAS[13] have demonstrated considerable success in identifying proteinogenic substrates. However, since these residues were defined based on a single co-crystal structure from one species and substrate class, their generalizability across phylogenetically diverse A-domains remains limited. Moreover, the scarcity of nonproteinogenic substrates dataset has significantly hindered progress in substrate specificity prediction. Therefore, it is advantageous to redefine SCCs and ABP residues by incorporating a broader set of co-crystal structures representing diverse A-domains and substrates for more accurate and generalizable identification.

Current extraction of binding pocket residues in A-domains mainly relies on sequence alignment, but accuracy drops for domains distant from the GrsA-Phe structure or those recognizing non-proteinogenic substrates[3]. Additionally, key patterns are often unclear in 1D sequences but conserved in 3D structures[14]. To improve substrate specificity prediction, it is desirable to develop computational frameworks that incorporate structural information for more accurate identification of SCCs or ABP residues. Recent advances in deep learning have transformed computational biology by enabling the extraction of complex patterns from large-scale datasets, including implicit structural features[15]. Protein language models (PLMs) and chemical language models (CLMs), such as Evolutionary Scale Modeling 2 (ESM2) and MoLFormer, exemplify this progress, leveraging self-supervised learning to uncover evolutionary and structural patterns from vast protein sequence and chemical structure corpora[16]. Transfer learning can further enhance these models by enabling the extraction of fine-grained structural and functional features, facilitating tasks such as protein function prediction and residue-level mutational impact analysis[15c, 17]. For instance, the ESM2-650M model has demonstrated accurate residue-residue contact prediction in both active and inactive peptides which directly facilitates the engineering of antimicrobial peptides[15c]. More recently, protein structures and the spatial relationships between residues have been effectively represented as graphs for analysis using graph neural networks (GNNs)[18]. By reframing binding pocket residue identification from a sequence-based to a graph-based problem, it becomes possible to harness the conserved spatial patterns characteristic of A-domains. This shift holds significant potential for improving the precision of ABP residues identification and advancing substrate specificity prediction.

Another challenge in substrate specificity prediction lies in the probabilistic formulation of prediction models. Most existing tools rely on softmax-based multiclass classification, which treats substrate labels as mutually exclusive and redistributes probabilities across all candidate substrates. This coupling can distort confidence estimates for individual substrates. Accordingly, there is growing interest in alternative formulations that assign independent confidence scores to each candidate substrate. Recent progress in protein-ligand interaction prediction provides a useful conceptual and technical foundation for such developments. Features derived from PLMs and CLMs can be leveraged to construct high-dimensional representations of binding pockets and ligands, and contrastive learning has emerged as an effective strategy to align these representations in a shared embedding space[19]. In this framework, contrastive objectives pull together embeddings of true protein-ligand pairs while pushing apart mismatched pairs, thereby enhancing representation quality and enabling more accurate interaction scoring and ranking. Emerging alignment-free approaches, inspired by virtual screening, predict protein–substrate interactions directly from protein sequences and molecular descriptors such as SMILES strings. State-of-the-art models trained on binding-affinity datasets enable systematic identification of likely interaction pairs, while architectures that integrate PLMs and CLMs offer improved robustness and adaptability to unseen substrates[20].

To address these limitations, we redefined the A-domain binding pocket as 27 residues (27-AA ABP) derived from evolutionarily diverse pocket–substrate pairs, forming a compact, spatially coherent cavity that preserves critical contacts, reduces structural noise, and can be efficiently encoded by a graph attention network. Building on this, we developed DeepAden, a two-stage ensemble tool for A-domain substrate specificity prediction, composed of three innovative modules: (1) a residual neural network[21] (ResNet)-based residue contact predictor that utilizes spatial constraints derived from PLM attention maps to capture the structural features of A-domains; (2) a graph attention network-based ABP prediction module (ABP-GAT) that combines residue-level features with probabilistic contact edges, enabling dynamic modeling of binding pocket variation without reliance on predefined templates; and (3) a contrastive learning framework that aligns cross-modal representations of ABPs and substrates, while addressing class imbalance through SHapley Additive exPlanations (SHAP)[22]-guided data augmentation. In real-world accuracy comparisons, DeepAden achieves performance that is comparable to or better than existing A-domain substrate prediction tools, particularly under low-homology and nonproteinogenic settings, in the examples presented in this work. Additionally, we applied DeepAden to predict A-domain substrate specificities within orphan NRPS BGCs and link them to candidate metabolites, followed by in vitro experimental validation in representative case studies. Our work establishes DeepAden as a reliable tool for A-domain substrate specificity prediction, highlighting the potential of AI to accelerate natural product discovery.

## 2. Results

### 2.1. Overview of DeepAden

To rapidly identify the residues within the pocket involved in substrate recognition of A-domains, we first developed a module named ABP-GAT, which integrated ESM2 with a graph attention network. ABP-GAT was designed to locate potential substrate-binding pockets using only the amino acid sequence, without relying on explicit structural data. It first transformed each sequence into a protein graph, where individual residues served as nodes. Each node combined two types of information, namely the sequence embeddings learned by ESM2 and the physicochemical properties of each residue. To determine how residues were connected within this graph, we employed a residual neural network (ResNet) that took the attention map from ESM2 as input to predict residue-residue contacts (Supplementary File). The predicted contact map defined the edges between residues, thereby establishing the overall graph structure. Once the graph was constructed, ABP-GAT was trained to identify which residues were most likely to form the substrate-binding pocket (Figure 1A).

**Figure 1.**
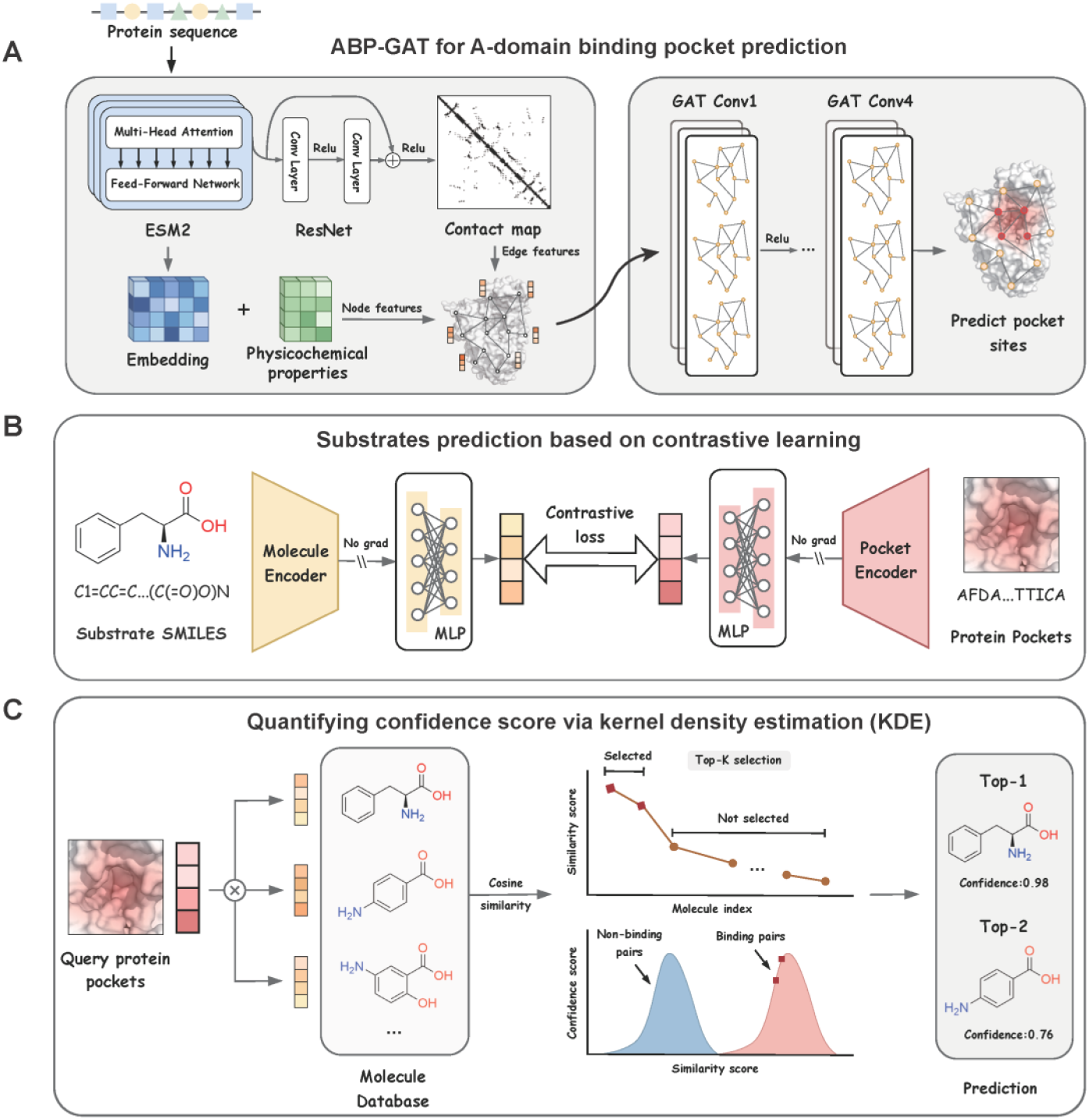
Overview of DeepAden. **(A)** A-domain binding pocket prediction with ABP-GAT. The protein sequence is encoded by ESM2, and residue embeddings are concatenated with physicochemical features to form node representations. A residue– residue contact map, predicted from ESM2 features by a small ResNet-based module (Methods), defines graph edges. A graph attention network (GAT) assigns pocket-forming probabilities to residues and outputs the predicted substrate-binding pocket. **(B)** Substrate prediction by contrastive learning. A pocket encoder (ABP-ESM2) and a molecule encoder (MoLFormer) produce embeddings that are further projected by trainable MLP heads. The model pulls embeddings of true pocket–substrate binding pairs together and pushes apart non-binding pairs in a shared embedding space. **(C)** Confidence estimation using kernel density estimation (KDE). Separate KDE are fitted to the cosine similarities of true binding pairs (positive distribution) and non-binding pairs (negative distribution) from training data. The similarity of a new pocket-substrate pair is evaluated under this KDE: scores in high-density regions are assigned higher confidence, whereas low-density scores are considered less reliable.

After identifying the binding-pocket residues, DeepAden predicted substrate specificity by matching each pocket with potential substrates. This task was formulated as a retrieval problem, where the model learned to pair each pocket with its true substrate among many candidates. To achieve this, we used two pretrained encoders: A fine-tuned ESM2 for ABP feature representation (ABP-ESM2) for pocket sequences and MoLFormer for substrate SMILES strings (See Methods). Their outputs were passed through trainable multilayer perceptron (MLP) projection heads, which were optimized so that embeddings of interacting pocket-substrate pairs were pulled closer together, whereas embeddings of non-interacting pairs were pushed farther apart in the shared embedding space (Figure 1B). To further enhance generalization, we trained an independent pairwise random forest classifier on pocket-substrate features and used SHAP to identify key residues in the A-domain pocket. Residues with high SHAP importance were kept fixed, whereas the three positions with the lowest SHAP importance in each pocket were selectively mutated using BLOSUM-62 substitutions. This SHAP-guided augmentation generated new, biologically plausible pocket-substrate pairs and effectively expanded the training space for DeepAden.

Finally, we quantified the confidence of each prediction by modeling the distribution of cosine similarity scores between pocket and substrate embeddings using kernel density estimation[23]. By obtaining a smooth, non-parametric estimate of the similarity-score density, we identified score regions corresponding to densely populated, reliable matches versus sparse, ambiguous ones, and used these density-based thresholds as a confidence criterion to distinguish high-confidence substrate matches from uncertain cases (Figure 1C). To train an evaluate this framework, we curated datasets from two recent studies, NRPStransformer (4,430 entries) and PARAS (3,257 entries), and further expanded them by manual literature search, yielding a final training set of 4,545 A-domain sequences from the bacterial phylum, fungal kingdom, metazoan kingdom, and plant kingdom (Figure S1). The detailed criteria and procedures for data curation, filtering, and cleaning are described in the Supplementary file.

### 2.2. DeepAden achieves robust binding pocket prediction across evolutionary divergence via GAT

The traditional coding approaches such as 10-AA[6a] and its derived 13-AA[24] and 17-AA[25] specificity-conferring codes (SCCs) face limitations in terms of predicting diverse substrates due to evolutionary divergence in A-domain structures. The 34-AA A-domain binding pocket (ABP) represented a major advancement by incorporating structural elements within an 8 Å substrate radius in GrsA-Phe (PDB: 1amu)[6b]. However, SCC and ABP definitions derived from this single GrsA-Phe cocrystal structure show limited generalizability across phylogenetically diverse A-domains and nonproteinogenic substrates. To obtain a more broadly applicable and structurally grounded ABP definition, we carried out a comprehensive structural analysis of 10 available A-domain-substrate cocrystal structures that contain bound amino-acid-like ligands without AMP, including both proteinogenic and nonproteinogenic substrates (Figure 2A). Noncovalent interactions that mediate substrate recognition, such as hydrogen bonds, salt bridges, and van der Waals contacts, typically operate within ∼6 Å[26], and several recent protein-ligand modeling frameworks (e.g., DrugCLIP[16b]) have adopted a 6 Å ligand-centered radius to define pocket regions. Guided by this biophysical rationale, we systematically compared pocket definitions based on residues within either a 6 Å or a traditional 8 Å radius of the bound substrate across these 10 cocrystal complexes. By mapping all 10 structures to a common sequence-structure alignment, we defined two ABP sets as the union across all structures of contacting residues under each distance cutoff: 27-AA ABP comprising all positions that lie within 6 Å of the substrate (Figure 2A), and 49-AA ABP comprising all positions that lie within 8 Å (Table S1). These two sets provide directly comparable pocket definitions derived from the same structural corpus but using different distance thresholds.

**Figure 2.**
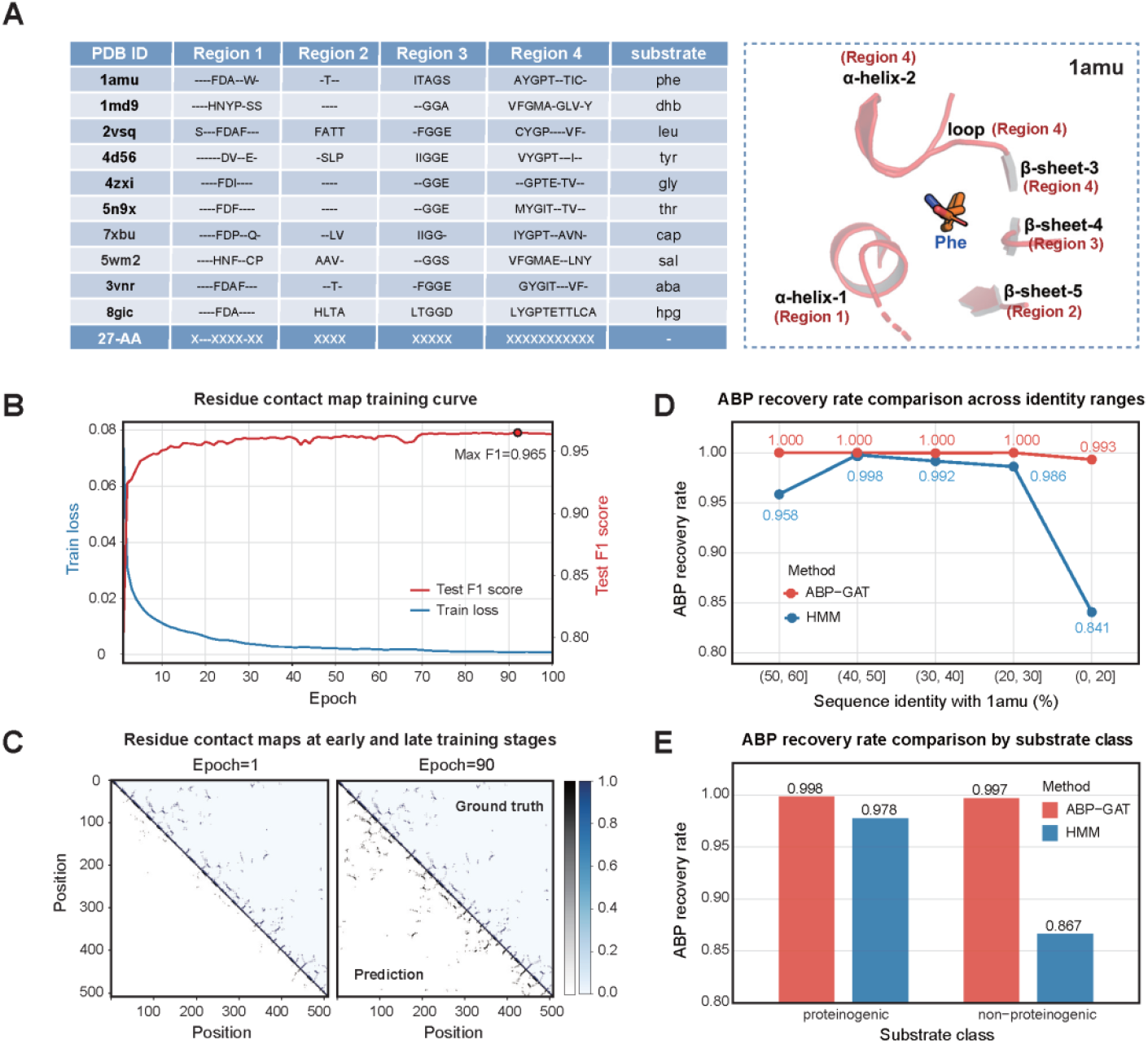
Performance of ABP-GAT model for binding pocket prediction. **(A)** ten A-domain cocrystal structures with bound substrates are used to define a canonical 27-residue A-domain binding pocket (27-AAABP). Pocket residues are defined as those within 6 Å of the substrate and are distributed across four conserved secondary structure regions. **(B)** A ResNet-based contact-map predictor trained on 78 A-domain crystal structures reach a peak F1-score of 0.965 on the test set, showing accurate learning of residue–residue contacts. **(C)** Predicted contact maps (lower triangles) at early and late training stages progressively approach the experimental contact maps (upper triangles), illustrating that the predictor captures the underlying structural constraints. **(D)** For 4,545 A-domain sequences, ABP-GAT (red) and a 1amu-based HMM method (blue) is evaluated for recovery of the complete 27-AAABP. Sequences are binned by their identity to the reference structure 1amu, and ABP-GAT maintains high recovery across all identity ranges. **(E)** ABP-GAT and the HMM method are further compared on A-domains recognizing proteinogenic and non-proteinogenic substrates. ABP-GAT maintains high ABP recovery in both classes, while the HMM performs substantially worse on non-proteinogenic substrates.

To overcome the limitations of HMM algorithms, which rely exclusively on primary sequence alignments for binding-pocket prediction, we next exploited both sequence and structural information from 78 published A-domain crystal structures to train a ResNet-based residue contact map predictor (Figure 2B). This model achieved an F1 score of 0.965 on the test dataset and accurately captured residue-level interaction patterns within A-domains, as illustrated by the convergence of predicted and experimental contact maps over training (Figure 2C), providing residue-level structural constraints for downstream ABP prediction. Using these predicted contact maps as edge information and protein language model embeddings as node features, we developed the ABP-GAT model for residue-level pocket prediction, which achieved a per-residue F1 score of 0.979 on the test dataset. To quantitatively assess how the distance cutoff affects pocket learnability, we trained ABP-GAT under both the 6 Å-based 27-AA and the 8 Å-based 49-AA definitions using the same training settings, and evaluated performance at the ABP-set level, defined as whether the entire ABP was correctly recovered as a complete set of residues for a given A-domain. Under the 27-AA definition, ABP-GAT correctly recovered the full pocket for 0.9980 of A-domains, whereas under the 49-AA definition this fraction decreased to 0.9386, indicating that the 27-AA is more precisely and consistently recoverable and is therefore better suited for subsequent modeling. Notably, all 27-AA are part of the 34-AA defined by Rausch et al[6b]. and extend the canonical 10-AA defined by Stachelhaus et al[6a]. (Table S2)

We next benchmarked ABP-GAT against an HMM-based method under the 6 Å-based 27-AA ABP definition. To this end, we evaluated a curated set of 4,545 A-domains that share 0–60% sequence identity with the reference A-domain GrsA-Phe (1amu) and cover diverse taxonomic origins. For the HMM baseline, we used the NRPSPredictor2 profile built on 1amu to infer binding-pocket residues. For both methods, pocket prediction accuracy was assessed at the ABP-set level, defined as whether the entire 27-AA was correctly recovered as a complete set of residues for a given A-domain. Across all sequence-identity bins within the 0–60% range, the ABP-set recovery rate of the HMM approach declined sharply as identity to 1amu decreased, whereas ABP-GAT maintained near-perfect accuracy in every bin (Figure 2D). We further stratified A-domains by substrate class and observed that ABP-GAT achieved similarly high recovery rates for both proteinogenic and nonproteinogenic substrates, while the HMM model performed substantially worse for nonproteinogenic substrates (Figure 2E). These results demonstrate that the learned graph-based model generalizes markedly better than the 1amu-based HMM profile, particularly for evolutionarily distant A-domains and those specific for nonproteinogenic substrates.

### 2.3. Conservative and functional residue recognition through evolutionary and SHAP analyses

Employing the ABP-GAT model with sequence-based inputs, we obtained 4,545 27-AA ABP sequences with known substrate annotations (labelled) and 186,758 27-AA ABP sequences without substrate annotations (unlabelled) from GTDB Release 225 and UniRef release 2024_06[27] (Supplementary File). To assess how strongly these short ABP segments encode substrate information, we constructed a phylogenetic tree based solely on the 27-AA ABP alignment (Figure 3A) and examined how this “pocket-level” phylogeny relates to organismal classification. Visual inspection of the tree revealed substrate-enriched sectors around the perimeter, for example, regions dominated by glycine, threonine, cysteine, alanine, serine, branched-chain hydrophobic residues (Val/Ile/Leu), and basic residues (Lys/Orn/Arg). Within these substrate-enriched regions, however, sequences from multiple bacterial phyla and from eukaryotes are intermingled, and long branches that are simultaneously dominated by a single phylum and a single substrate label are relatively uncommon. This pattern suggests that, at the level of the substrate-binding pocket, sequence similarity is shaped primarily by substrate-related evolutionary pressures and is only weakly aligned with broad organismal lineages. Notably, fungal ABPs (black ring) are scattered across multiple substrate-enriched sectors and frequently co-occur with bacterial ABPs that share the same substrate label (Figure 3A). We then quantified how tightly each substrate label is clustered on the 27-AA ABP tree using an Adjusted Rand Index (ARI)[28] analysis (Figure 3B). Substrates such as piperazic acid and the small or polar amino acids glycine, threonine, and cysteine exhibited high ARI values, indicating that their corresponding ABPs are concentrated within relatively compact regions of the tree. In contrast, substrates with lower ARI values are more widely dispersed across the phylogeny, consistent with more heterogeneous or promiscuous binding pockets.

**Figure 3.**
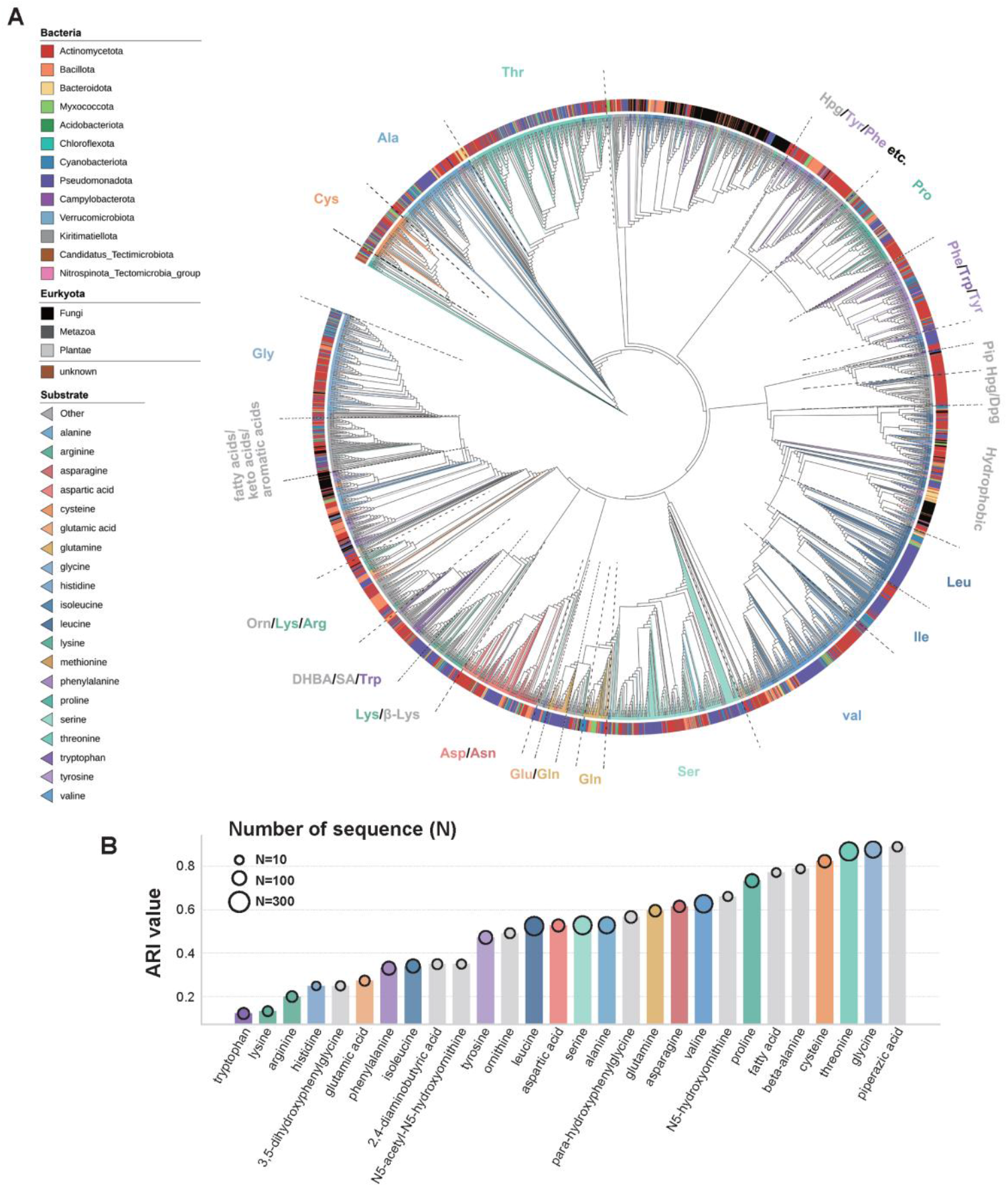
Evolutionary relationships and substrate specificity of A-domain binding pocket (ABP) sequences. **(A)** Circular phylogenetic tree constructed from a curated dataset of 4,545 ABP sequences. Each leaf represents a single A domain, with branches colored by substrate and the outer ring indicating taxonomic affiliation. **(B)** Bar plot showing the adjusted Rand index (ARI), quantifying how well clusters of bacterial ABP binding-pocket sequences in the phylogenetic tree agree with their substrate labels. The tree was cut at a fixed height (H = 2.0 substitutions) to define clusters, which were compared with substrate labels using ARI (1 = perfect agreement, ≈0 = random); point size indicates the number of sequences collected for each substrate label.

We observed that positions 2, 3, 15, 18, 19, and 22 are more highly conserved among labelled pockets (Figure 4A), indicating a limited set of tolerated amino acids at these sites. Such conservation suggests that these positions contribute to maintaining the pocket architecture and/or to the recognition of particular substrate types. Because substrate annotations in our dataset are strongly imbalanced, further analyses are needed to clarify the substrate-specific contributions of individual positions. Recently, explainable artificial intelligence methods such as SHAP have emerged to decode model decisions by quantifying the contributions of individual amino acids to protein properties[29]. To move from these global patterns to a residue-resolved view of substrate recognition, we next quantified the contribution of each ABP position using model-agnostic feature attribution. Specifically, we leveraged SHAP to interpret predictions from machine-learning models trained on ABP sequences. Building on this approach, we fine-tuned the ESM2-650M model on a curated dataset of 191,303 ABP sequences (labelled and unlabelled), yielding a model referred to as ABP-ESM2 (See Methods). This model generated context-aware positional embeddings that exhibited clearer class separation in embedding space than the original ESM2 model, while effectively preserving positional information critical for downstream analyses. For the labelled subset of 4,545 ABP sequences, we applied an oversampling strategy combined with 5-fold cross-validation to train random-forest classifiers on pairwise substrate-versus-non-substrate discrimination tasks, fitting a separate classifier for each substrate class. SHAP values were then computed for each substrate-specific model directly on the ABP-ESM2 embeddings. Across all substrate classes, the SHAP analysis highlighted eight positions (1, 2, 3, 4, 9, 20, 24, and 27) with the largest overall contributions (Figure 4B and 4C). These positions cluster around non-carboxyl substrate moieties, side-chain interaction zones, and the only loop segment lining the pocket, indicating that contacts in these regions are a major source of discriminative information for substrate specificity. In particular, the loop-embedded positions 20 and 24 are located near the pocket entrance, suggesting that this flexible segment helps shape the local binding environment and modulate how different substrates are accommodated. The highly conserved positions 15, 19, and 22 show comparatively low SHAP contributions, suggesting that they provide little discriminative information for distinguishing substrates in our models and are more likely involved in maintaining the overall pocket scaffold than in fine-tuning substrate preference. Although positions 2 and 3 are highly conserved, they still help distinguish ABPs specific for amino-acid substrates from those recognizing non-amino-acid substrates (Figure S2). Together, these patterns indicate that combining conservation with SHAP-based feature attribution distinguishes structurally important pocket residues from those that directly encode substrate type.

**Figure 4.**
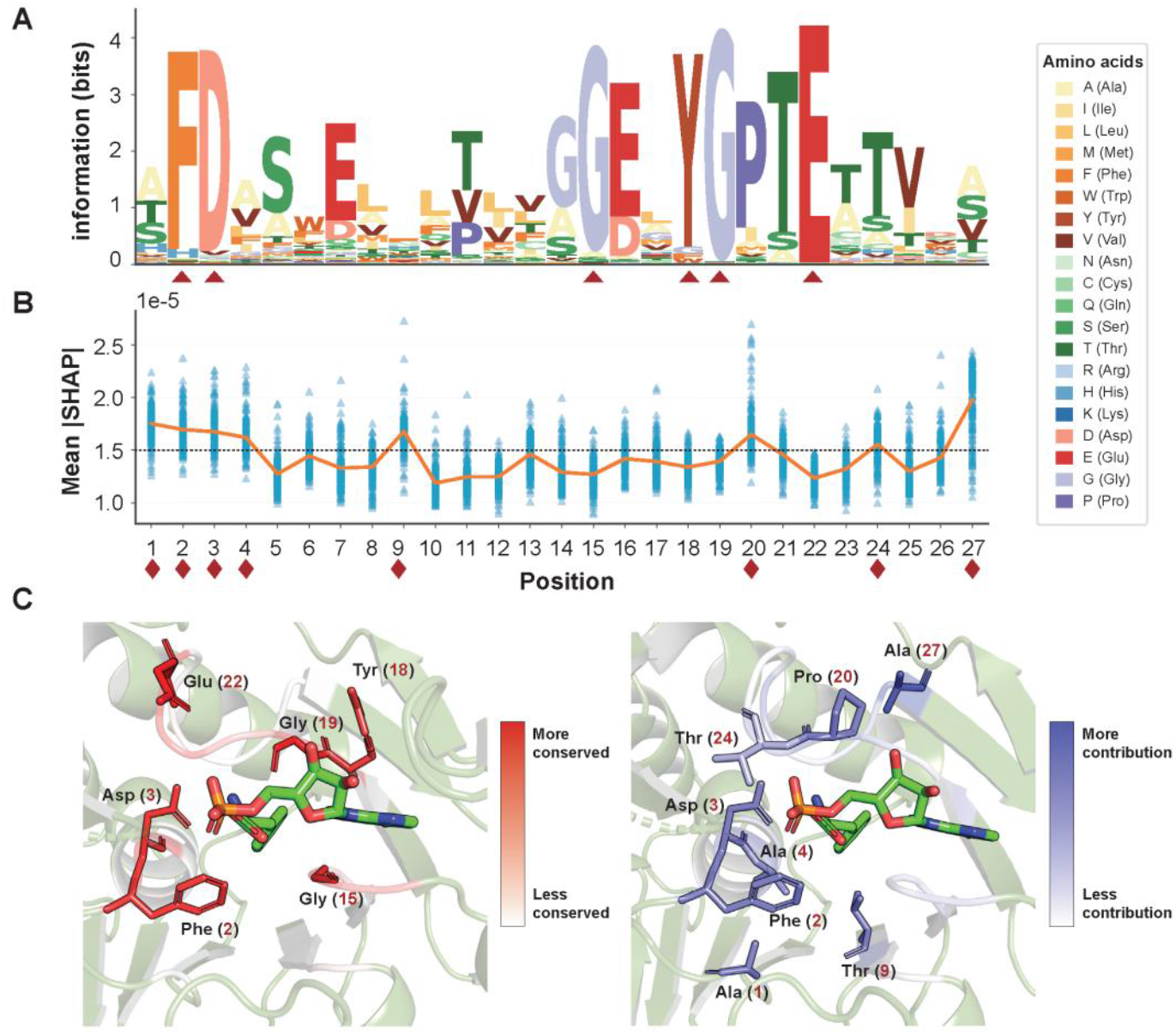
Conservation and substrate-specificity determinants in A-domain binding pockets (ABPs). **(A)** Sequence logo and conservation profile of 4,545 ABP sequences across the 27-AA binding-pocket positions. Triangles mark higher conserved residues. **(B)** Mean absolute SHAP value (|SHAP|) for each of the 27 ABP positions in pairwise random-forest classifiers distinguishing substrate classes, representing the contribution of each position to substrate specificity. Each point corresponds to the mean |SHAP| value for a single substrate class at the indicated position. Diamonds indicate positions with the higher contributions. **(C)** Conservation scores (left) and SHAP-based contribution scores (right) mapped onto the crystal structure of GrsA-PheA (PDB: 1AMU), highlighting residues that are both highly conserved and strongly associated with substrate specificity.

In DeepAden, SHAP-guided augmentation both guides position-specific substitutions for data augmentation and provides a framework for systematically identifying substrate-specific residues across characterized A-domains. In the 27-AA ABP phylogeny (Figure 3A), Cys-ABPs and Ser-ABPs form distinct clades, even though L-Cys and L-Ser differ only subtly in side-chain size and polarity. This observation prompted us to ask whether SHAP could resolve the fine-scale ABP differences that underlie this separation. For the clarity of the following description, the superscript numbers preceding the amino acid residues indicate their positions within the ABP sequence alignment results, whereas the numbers following the residues correspond to their positions in the resolved PDB structure. Class-specific SHAP heatmaps for Cys-ABPs and Ser-ABPs revealed pronounced differences, with position 20 showing the largest substrate-dependent shift in SHAP values and corresponding to ^20^Ala in Cys-ABPs and ^20^Pro in Ser-ABPs (Figure 5A). High-resolution structural comparisons between the Cys-AMP-bound (7en1) and Ser-AMP-bound (6ea3) complexes revealed distinct ligand coordination mechanisms consistent with these SHAP-highlighted differences (Figure 5C and F). In Ser-ABPs, ^20^Pro rigidifies the loop at the pocket entrance and packs against the ribose–phosphate moiety, restricting its conformational flexibility (Figure 5C and D). In Cys-ABPs, the corresponding ^20^Ala, together with neighboring ^18^Gly, creates a more flexible local environment that permits alternative orientations of the ribose-phosphate group (Figure 5E and F). This difference in loop flexibility is accompanied by a rearrangement of adenine interactions. In Cys-ABPs, the adenine ring adopts an “edge-to-face” π-π stacking interaction with Phe939 located outside the canonical binding pocket, tilting the ring upward and enabling two hydrogen bonds to form with ^17^Leu (Figure 5E and F). As a consequence, the ribose moiety is reoriented, with its O4′ atom positioned closer to the adenine ring than in the Ser-ABP complex. The increased local flexibility around positions 18 and 20 in Cys-ABPs also facilitates productive positioning of the Cys thiol group to hydrogen bond with ^19^Gly. Together, these case studies illustrate that SHAP-identified positions, particularly position 20, correspond to structurally meaningful interaction hotspots that differentiate Cys-ABPs and Ser-ABPs, thereby supporting both the biological plausibility of our attribution analysis and the rationale for constraining data augmentation to positions with low SHAP importance.

**Figure 5.**
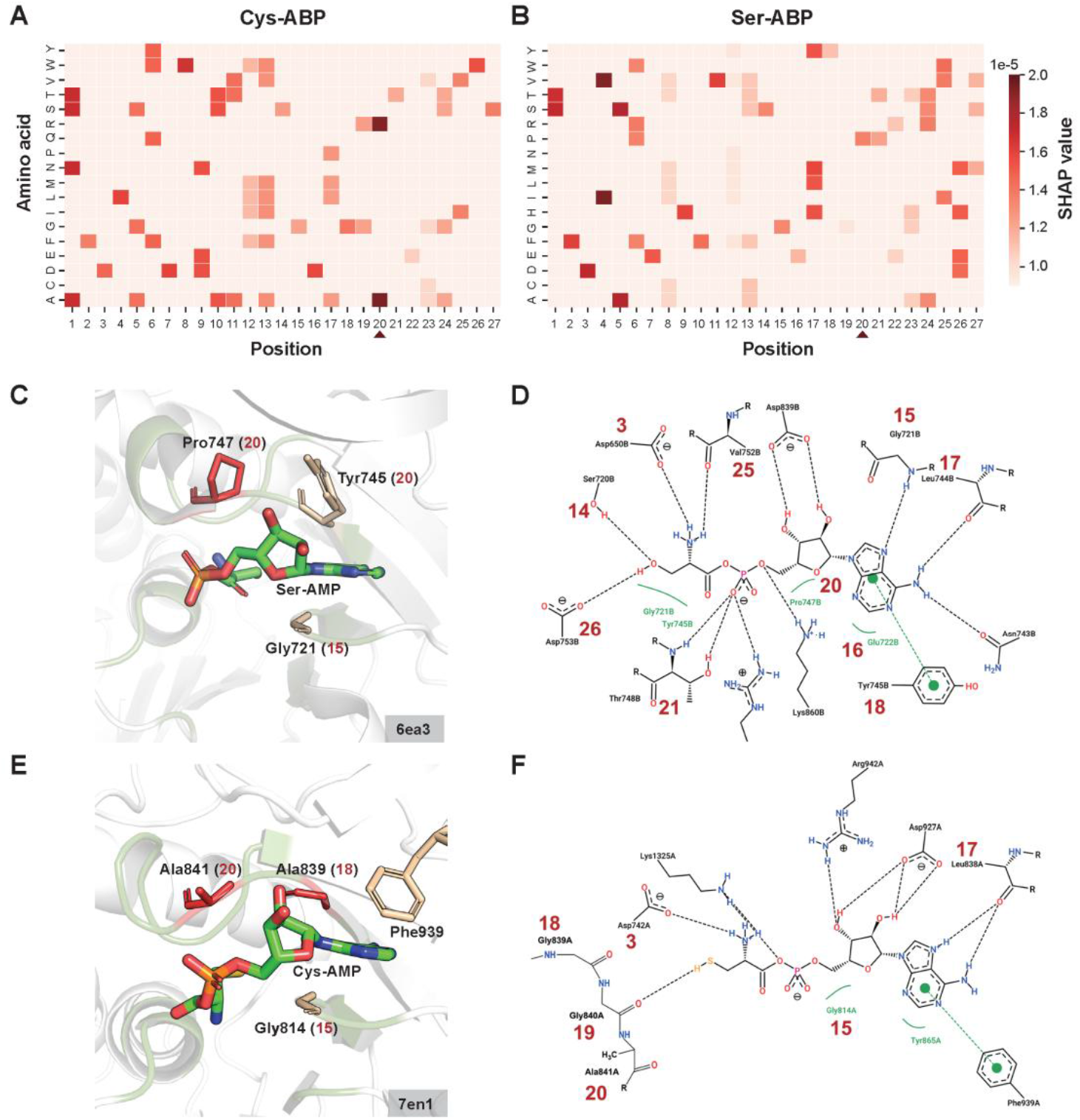
Structural and interaction analysis of functional residues in A-domain binding pocket (ABP) based on PchE-Cys and FscH-Ser crystal structures. **(A, B)** Heatmaps of positive SHAP values highlighting amino-acid contributions at each binding-pocket position for Cys-ABPs and Ser-ABPs, respectively, for the sequence clusters shown in Figure 3A. **(C)** Structural representation of contributing residues within the ABP of FscH-Ser (PDB: 6ea3). The bound ligand is Ser-AMP, containing an adenine ring and a pentose sugar. Red numbers in parentheses indicate positions in the 27-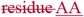 ABP, and black labels indicate amino acids and their positions in the protein sequence. **(D)** Two-dimensional interaction diagram of FscH-Ser (PDB: 6ea3). Interactions between protein and ligand are shown as dashed lines, with both interacting residues and the ligand drawn as structural formulas. Hydrophobic contacts are indicated by highlighted hydrophobic regions of the ligand and the labels of contacting residues, and dashed lines between aromatic rings denote π-π stacking interactions. **(E)** Structural representation of contributing residues within the ABP of PchE-Cys (PDB: 7en1). The bound ligand is Cys-AMP; residue annotations follow the same convention as in panel (C). **(F)** Two-dimensional interaction diagram of PchE-Cys (PDB: 7en1), with interaction types annotated as in panel (D).

### 2.4. DeepAden achieves SOTA performance in substrate specificity prediction including nonproteinogenic substrates

Current methods for predicting NRPS A-domain substrates perform poorly on recently identified nonproteinogenic substrates, largely because suitable training sequences are scarce and sequence alignment–based SCC extraction has clear limitations. To increase the effective data for these substrates, we augmented their ABP sequences. Simple oversampling or random perturbations risk overfitting and generate sequences with low biological plausibility, so we instead adopted a SHAP-guided augmentation strategy. Based on SHAP and conservation analyses (Figures 4 and 5), we distinguished residues that were evolutionarily conserved, functionally important, or located in interaction hotspots from residues that were weakly conserved, have minimal impact on model predictions, and lie outside structurally validated binding regions. Only the latter positions were selected for in silico mutagenesis, as substitutions at these sites are more likely to be tolerated without altering substrate specificity. Guided by observed substitution patterns and physicochemical similarity, we introduced small, conservative mutations at these low-influence positions, thereby expanding the local sequence space around under-represented nonproteinogenic substrates while preserving key recognition features (Figure S3). The augmented dataset was then used in the subsequent contrastive learning stage to enhance the model’s discriminative ability.

In the second stage, we trained the substrate prediction model within a contrastive learning framework using the augmented dataset. The data were split into training and validation sets at an 8:2 ratio, and 5-fold cross-validation was used for model training and hyperparameter tuning. After identifying the optimal hyperparameters, we retrained the model on the full dataset to obtain the final weights. The trained DeepAden model captured the characteristic interaction patterns between the A-domain binding pockets and substrate molecules, effectively distinguishing positive sample pairs (pocket–substrate pairs) from negative sample pairs (pocket–non-substrate pairs) (Figure 6A). In the embedding space, contrastive learning successfully shortened the distance between substrate molecules and their corresponding pockets (Figure S4). We next compared models trained on the original and augmented datasets. In 5-fold cross-validation, mean validation accuracy increased from 78.02%±1.58% with the original dataset to 90.57%±0.57% with the augmented dataset (p-value < 1e-4) (Figure 6B), indicating that SHAP-guided augmentation substantially improved the learned representations.

**Figure 6.**
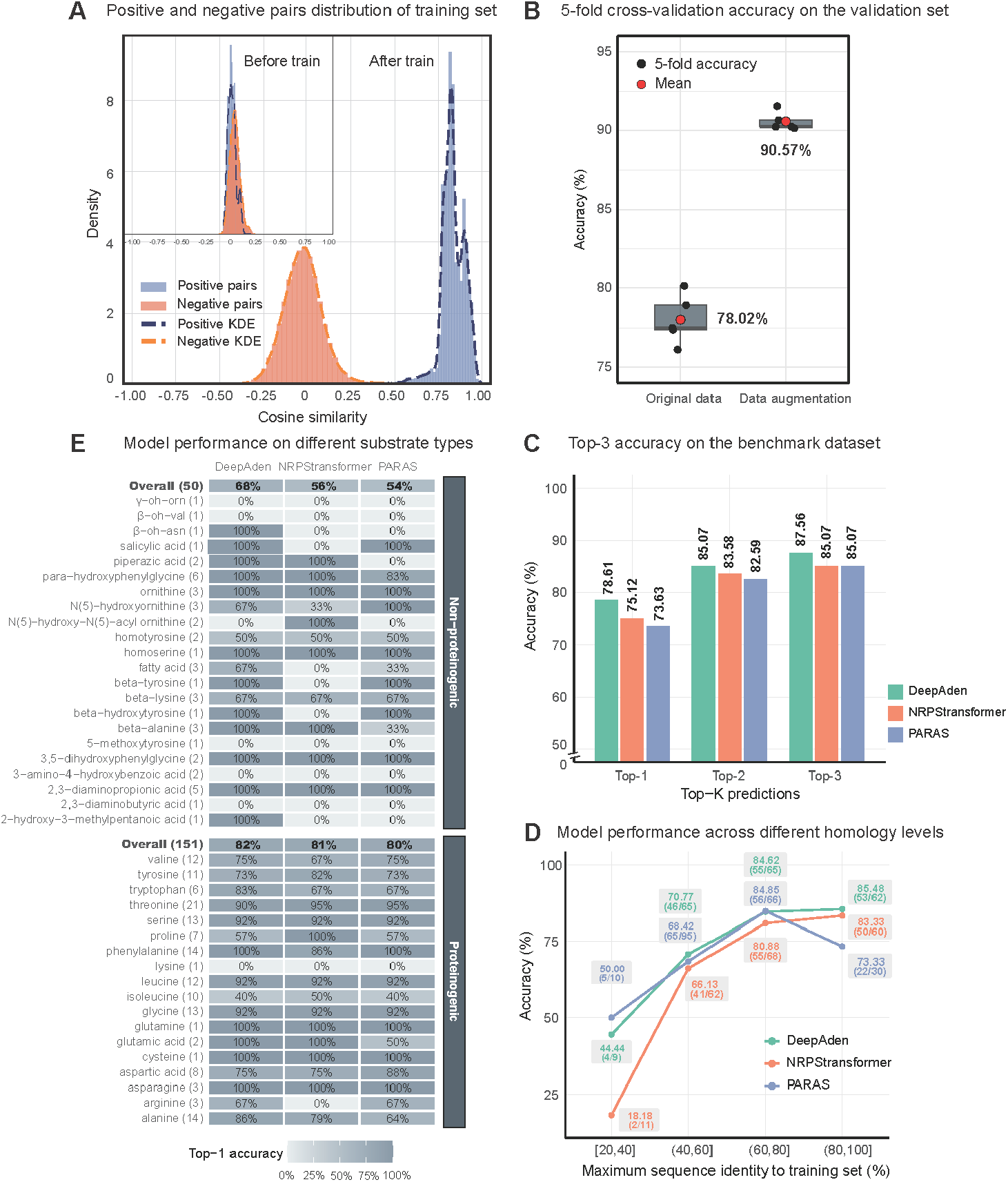
Model performance evaluation of DeepAden. **(A)** Cosine similarity distributions of positive (true pocket-substrate) and negative pairs before and after contrastive learning. After training, positive and negative pairs form two well-separated modes. **(B)** 5-fold cross-validation accuracy on the validation set increases from 78.02% ±1.58% on the original data to 90.57%±0.57% with SHAP-guided data augmentation, indicating that this strategy markedly improves representation learning for pocket-substrate interactions. **(C)** On an independent benchmark dataset of 201 A-domains, DeepAden attains slightly higher Top-1, Top-2, and Top-3 accuracy than NRPStransformer and PARAS. **(D)** Benchmark dataset were grouped according to their maximum sequence identity to the training set of each model. Accuracy decreases as sequence identity drops, but DeepAden maintains competitive performance across all homology levels. (E) For proteinogenic substrates (151 samples), all models performed similarly. For nonproteinogenic substrates (50 samples), DeepAden outperformed NRPStransformer by 12% and PARAS by 14%, demonstrating improved recognition of complex or rare substrates.

We next benchmarked DeepAden against two recently developed A-domain substrate prediction tools, NRPStransformer[30] and PARAS[13]. The benchmark set contained 201 A-domain entries with no overlap with the training data of the assessed tools. Of these, 155 sequences were taken from the NRPStransformer benchmark, and 46 sequences were manually curated from literature corresponding to newly added MIBiG 4.0 entries[31] (Supplementary File). On the 201-sequence benchmark, DeepAden achieved slightly higher top-3 accuracy than NRPStransformer and PARAS by 2.5–5%, although all three tools remained competitive and differences were not statistically significant in all subranges. None of the tools exceeded 80% in top-1 accuracy, but all showed substantial gains in top-2 accuracy, ranging from 6.5% to 9% (Figure 6C).

To assess the generalization capacity of each tool, particularly when handling A-domain sequences with low homology to their respective training sets, we calculated, for each of the 201 test sequences, its highest sequence identity to the training sequences of each tool and used this value as its similarity to that tool’s training data. Performance was then evaluated across different sequence identity intervals. DeepAden maintained competitive performance across multiple ranges. Specifically, in the high-similarity range (80, 100], DeepAden achieved the highest accuracy of 85.48% (53/62), with NRPStransformer performing comparably well. The lower performance of PARAS in this range may reflect differences in its training data coverage. In the moderate-similarity range (40, 80], the performance of all three tools was comparable, though accuracy decreased relative to the high-similarity interval. In the low-similarity range, predictive performance declined sharply for all models. Here, PARAS achieved 50.0% accuracy (5/10), followed by DeepAden at 44.44% (4/9), indicating that both retain some predictive capacity, whereas NRPStransformer reached only 18.18% (2/11) (Figure 6D). However, the small number of test sequences in this interval limits the statistical power of these comparisons. Across all tools, accuracy decreased with lower sequence identity to the training data, underscoring the importance of diverse training sets for robust substrate prediction in realistic, heterogeneous sequence spaces. Finally, we evaluated the ability of the three tools to predict two broad substrate categories: proteinogenic and nonproteinogenic amino acids. Overall, all tools performed similarly in predicting proteinogenic substrates. However, for non-proteinogenic substrates, DeepAden demonstrated superior predictive performance, outperforming NRPStransformer by 12% and PARAS by 14%, as illustrated in the heatmap (Figure 6E).

### 2.5. DeepAden localizes orphan NRPS BGCs and identifies their A-domains

As a proof of concept for DeepAden-assisted, structure-guided genome mining, we set out to link NRPS BGCs in the rice endophytic actinomycete *Streptomyces hygroscopicus* OsiSh-2 (GenBank accession JBPQZZ000000000) to their corresponding metabolites[32]. We first performed untargeted LC-MS/MS-based metabolomic analysis of culture extracts of *S. hygroscopicus* OsiSh-2. By matching accurate precursor *m/z* values, inferred molecular formulas, and diagnostic MS/MS fragments to literature data and entries in NPAtlas, Reaxys, and SciFinder, we putatively annotated two NRP families as nyuzenamide B and D and octaminomycin A-D. For the nyuzenamide family, the MS/MS spectra showed limited backbone fragmentation, consistent with previous reports on bicyclic peptides[33], so our annotation primarily relied on accurate mass, molecular formula, and high spectral similarity to published nyuzenamide reference spectra, supported by the similarity between the candidate BGC and previously reported nyuzenamide clusters (Figure S5). In contrast, the octaminomycin family displayed rich fragment ion coverage, which enabled a more detailed interpretation of the peptide backbone directly from the MS/MS data and subsequent comparison with A-domain substrate predictions (Figure S6).

The BGCs of *S. hygroscopicus OsiSh 2* were analyzed using antiSMASH v7.0, which identified 12 NRPS/NRPS-like BGCs (Figure S7). To establish gene cluster-compound correlations, we adopted an an A-domain–based strategy that proceeds from metabolite to genome and back to biosynthetic logic. First, we deduced the amino acid building blocks of the two target families from their putative structures inferred from the MS/MS data: nyuzenamides with the core peptide Phe-Pro-Leu-Gly-Trp-Asn-Thr-Hpg-Val-Gly, and octaminomycins with the core peptide Thr-Phe-Leu-Val-Pro-Leu-Tyr(Phe)-Pro(Pip). Notably, given the potential involvement of various modifying enzymes (e.g. isomerases and methyltransferases) during biosynthesis, we focused exclusively on the core scaffold derived from the initially recognized A-domain substrates, without accounting for late-stage tailoring modifications that may generate derivative structures. DeepAden was then used to predict substrate specificities for all A-domains encoded in the candidate NRPS BGCs. For each A-domain, DeepAden independently computed a score for every candidate substrate, allowing multiple plausible substrates to be considered for a single domain when appropriate. By systematically aligning the DeepAden substrate profiles with the amino acid compositions of the target core peptides, we were able to assign one NRPS BGC as the putative nyuzenamide cluster and another as the putative octaminomycin cluster. The GBK files of these two BGCs have been deposited in our GitHub repository.

For the nyuzenamide BGC, DeepAden recovered the expected substrate preference at every position of the core peptide scaffold (Figure 7A, B), thereby supporting a consistent cluster–product assignment at the level of the peptide backbone. In the case of the octaminomycin BGC, the A-domain in module M22 was predicted to incorporate Phe, but with a low confidence score (0.24), whereas the true substrate Leu ranked second with a score of 0.15. Notably, the A-domains in modules M23 and M24 showed recognition of multiple substrates (M23: Tyr/Phe; M24: Pro/Pip). In both modules, the primary substrates (Tyr and Pro) ranked first with confidence scores of 1.0, while the alternative substrates were still captured among the top-ranked predictions but with substantially lower scores (Phe ranked second with 0.18 in M23, and Pip ranked third with 0.02 in M24) (Figure 7C, D). This pattern indicates that the presence of these alternative amino acids is recognized by the model, albeit with markedly lower confidence than the dominant assignments.

**Figure 7.**
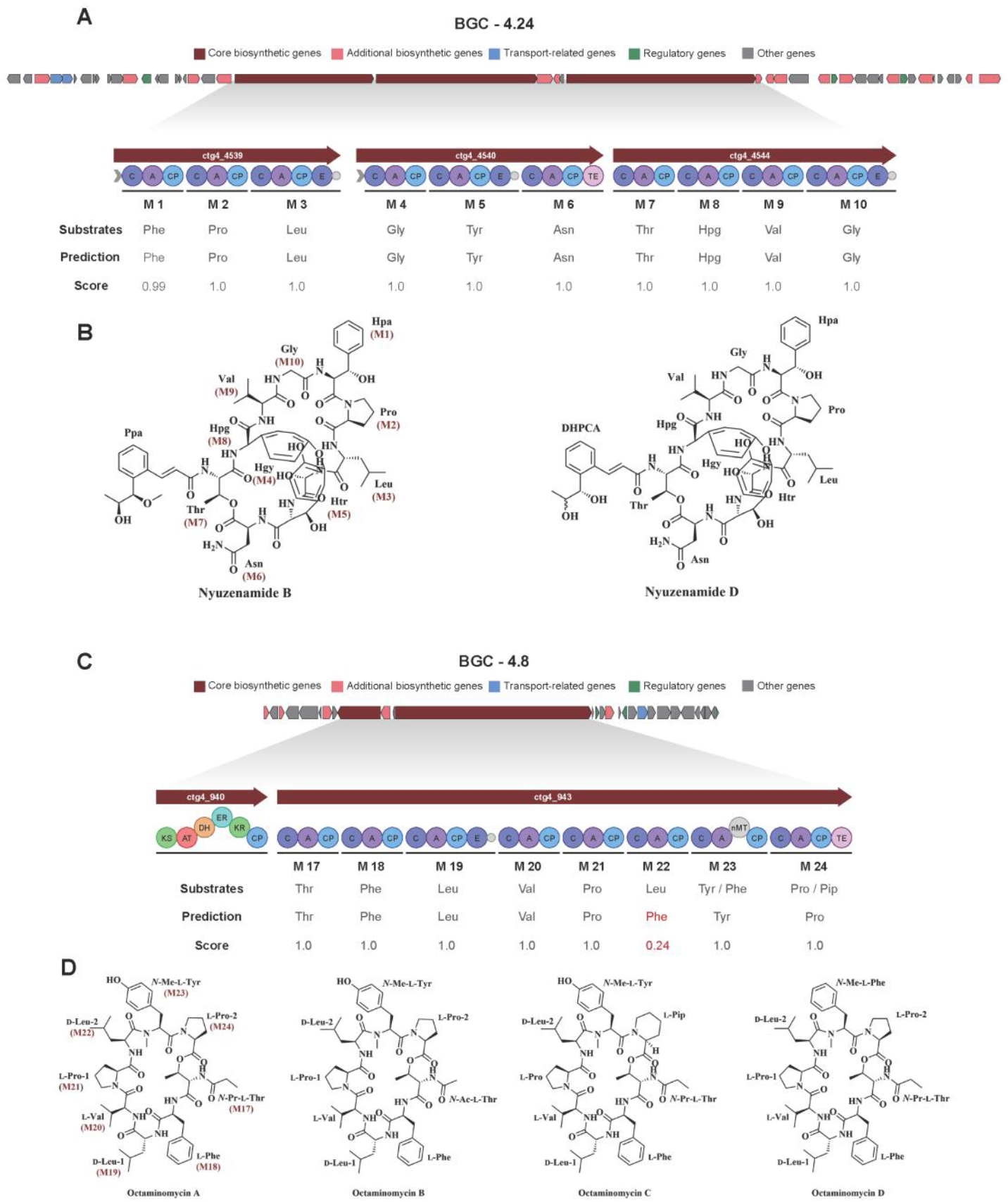
DeepAden-guided, structure-based assignment of orphan NRPS BGCs in *Streptomyces hygroscopicus* OsiSh-2. **(A)** Genomic context and NRPS organization of BGC-4.24 with DeepAden substrate predictions and scores for each A-domain (M1-M9). **(B)** Nyuzenamide B and D structures with their core peptide (Phe–Pro–Leu–Gly–Trp–Asn–Thr–Hpg– Val–Gly) mapped to the corresponding A-domains in BGC-4.24, supporting its assignment as the nyuzenamide BGC. **(C)** Genomic context and NRPS organization of BGC-4.8 with DeepAden substrate predictions and scores for A-domains M17-M24, including positions with experimentally observed alternative substrates (M23: Tyr/Phe; M24: Pro/Pip). **(D)** Octaminomycins A–D with the core peptide (Thr–Phe–Leu–Val–Pro–Leu–Tyr(Phe)–Pro(Pip)) mapped to BGC-4.8. DeepAden recovers the primary substrates and ranks the alternative substrates among the top predicted candidates. Detailed MS/MS-based structural inferences are provided in Figure S5 and S6.

## 3. Discussion

Substrate adenylation by A-domains is essentially a protein-ligand interaction process, requiring precise localization of the binding pocket and accurate substrate pairing for reliable prediction. With the development of deep learning, an increasing number of algorithmic frameworks have been proposed to tackle such interaction prediction tasks, particularly those involving high-dimensional and structured biological data[34]. Recent advances in pretrained PLMs have enabled the integration of structural context into A-domain representation learning. Meanwhile, contrastive learning has proven effective in protein-ligand interaction prediction[19]. In this study, we propose DeepAden, a two-stage framework that first trains a graph attention network to identify the binding pocket surrounding the target substrate by fusing sequence features with structural embeddings derived from a protein language model. In the second stage, it employs cross-modal contrastive learning to jointly encode pocket-substrate interactions using both protein and chemical language models, effectively addressing challenges arising from evolutionary divergence in A-domain substrate prediction. By integrating all these modules, DeepAden not only provides insights into substrate binding but also enables effective substrate screening using only A-domain sequences, without requiring any 3D structural data. This eliminates the dependence on 3D data and paves the way for robust learning across large-scale sequence databases. To support the natural product research community, we developed a user-friendly web server for ABP residue identification and substrate prediction

Recent methods have started to derived A-domain signatures by aligning structures predicted by ColabFold[13], but such approaches remain difficult to scale to genome-level analyses and are sensitive to structural prediction uncertainties. At the same time, sequence-alignment–based ABP extraction degrades markedly on evolutionarily divergent A-domains[3], limiting its applicability to nonredundant NRPS repertoires. To address these issues, we developed ABP-GAT, which mitigates the impact of evolutionary divergence by inferring ABPs directly from local pocket environments and maintains robust performance on nonhomologous A-domains with only 30-40% sequence identity to 1amu. Our analyses further show that a compact 27-AA pocket defined by a 6 Å cutoff is sufficient to support accurate substrate prediction. This 27-AA pocket achieves substrate prediction performance comparable to two state-of-the-art tools, NRPStransformer[30] (which uses the flavodoxin-like subdomain) and PARAS (which uses a 34-AA pocket)[13], indicating that a 27-AA pocket captures enough local information for reliable substrate inference. Moreover, when we stratify performance by global sequence identity, the pocket-based models (DeepAden and PARAS) exhibit reduced sensitivity to global homology compared with the full-length-based NRPStransformer, consistent with their underlying design principles. The ABP extraction method we developed demonstrates broad applicability and may provide a useful tool for further research. Building on this approach, future studies can investigate the functional mechanisms of these residues across diverse A-domains, which may help improve the accuracy and applicability of substrate-specificity prediction algorithms. To support broader usage, we offer predicted ABPs through both our command-line tools and web server, allowing users to explore the diversity and evolutionary dynamics of pocket sequences. It is worth noting that while our 27-AA pocket within a 6 Å universal boundary balances computational efficiency and generalization, its static boundaries may be limited in their ability to recognize large substrates and accommodate protein conformational changes[35]. Future research should explore dynamic pocket delineation strategies that incorporate molecular docking energy landscapes or residue-substrate contact frequency analyses. Such adaptive methods could enhance the accuracy of binding-interface characterization and improve substrate-specificity prediction performance.

DeepAden employs a contrastive learning-based independent modeling strategy that generates separate predictions and distinct confidence scores for each pocket–substrate pair. This mechanism reduces direct competition among substrates and is, in principle, more compatible with the multi-substrate nature of biochemical reactions at the A-domain level. In contrast, many existing tools utilize softmax functions to normalize probabilities across substrate libraries[10, 13]. Consequently, their mutual-exclusivity assumption can lead to averaged low-probability predictions or a single dominant substrate, which may under-represent potential multi-substrate usage. However, recent study have demonstrated that A-domain-level information alone does not fully determine whether a given amino acid can be incorporated into the final product, as additional constraints arise from the PCP domain and the broader module-level context[36]. This distinction is exemplified by our analyses of the nyuzenamide and octaminomycin BGCs (Figure 7), where DeepAden accurately recovers primary substrate preferences and surfaces experimentally validated alternative substrates within the top-3 predictions but assigns them lower confidence than the dominant amino acids. Consequently, although DeepAden’s independent-probability design is, in principle, capable of representing substrate promiscuity, its current predictions should not be taken as definitive evidence of true biochemical multi-substrate usage and should be complemented by PCP-domain analysis, module-level context, and targeted biochemical assays. Rather, lower-ranked or lower-confidence entries within the top k predictions are best treated as hypotheses that may reflect latent pocket-level promiscuity, but still require corroboration from PCP-domain features, module- and pathway-level context, or direct experimental validation. In practice, users interested in potential promiscuity should therefore cautiously inspect top-k predictions rather than relying solely on a single top-1 assignment, and interpret alternative substrates in light of structural and biochemical constraints downstream of the A-domain. To address these challenges, a promising future direction is to develop multimodal datasets that integrate atomic-level binding-pocket features and to design conformation feature extractors on the basis of geometric deep learning algorithms that enable “sequence-to-conformation-to-substrate” prediction[37].

Furthermore, it is important to acknowledge the inherent limitations of substrate-prediction tools in the broader context of natural product discovery. While DeepAden shows strong performance in predicting A-domain specificity on our benchmark datasets (Figure 6), the translation of these predictions into full compound structures is complicated by noncanonical NRPS assembly logic. Several NRPS BGCs do not follow the standard colinearity rule. For instance, the biosynthesis of surugamides[38], wollamides[39], and desotamides[40] involves iterative mechanisms, trans-acting domains, or module skipping, which decouple the linear gene sequence from the final chemical structure. In such cases, accurate substrate prediction is only one piece of the puzzle. Therefore, while DeepAden provides high-confidence predictions for monomer incorporation, these results must be integrated with detailed biosynthetic-logic analysis to infer plausible natural product scaffolds rather than to claim fully resolved structures. Notably, the 27-AA ABP proposed in this study provides a compact and informative representation of the binding pocket that can support such integrative analyses. Given the performance of DeepAden in predicting nonproteinogenic substrates that exhibit high evolutionary divergence from GrsA-Phe, we anticipate that this tool will facilitate substrate-level functional annotation and prioritization of NRPS BGCs identified by genome mining, thereby providing useful support for natural-product-based drug discovery.

## 4. Methods

DeepAden comprises two key components: an A-domain binding pocket prediction model and a substrate specificity prediction model. This section is organized into three main parts: “A GAT-based node classification model for A-domain binding pocket prediction”, “SHAP-guided functional residue recognition in APBs” and “Contrastive learning for pocket-substrate binding prediction”. The usage of webserver was supplied in supplementary file.

### 4.1. A GAT-based node classification model for A-domain binding pocket prediction

DeepAden processes 1D A-domain sequences into 2D graphs for candidate substrate binding pocket prediction. Binding pockets were predefined from 10 A-domain–substrate cocrystal complexes containing proteinogenic and nonproteinogenic ligands by selecting residues within a given distance of the bound substrates. The 78 A-domain crystal structures were used to train the GAT model, and for the remaining structures, the corresponding active pocket residues were determined through structural alignment using the align command in PyMOL v3.1.0. Details of data collection and preprocessing are provided in the Supplementary File. Using a 6 Å cutoff, we obtained a 27-residue pocket (27-AA), and using an 8 Å cutoff on the same structural set, we obtained a more inclusive 49-residue pocket (49-AA). Both 27-AA and 49-AA were trained in the following framework. The 27-AA-based ABP-GAT model was embedded in DeepAden.

#### 4.1.1. PLM-based graph construction

For a given 1D amino acid sequence with *N*_*p*_ residues, DeepAden generates a 2D protein contact map graph with *N*_*p*_ residue nodes using ESM2. The protein graph is characterized by 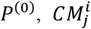 and 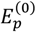, representing the residue node features, the contact map matrix that connects the nodes, and the edge features describing these connections, respectively. A residue feature is computed as follows:

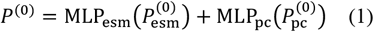

where 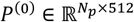 represents the computed residue feature; 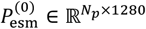 represents the residue-type feature obtained from the final layer of ESM2; and 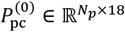 represents the physicochemical properties that encode the residue weights, pK values, the hydrophobicity status, and whether the residue is aliphatic, aromatic, neutral or charged. The contact map matrix 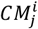 is predicted via ResNet with three residual blocks on the ESM2 attention map, and the edge features 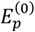 are obtained from the predicted contact probabilities obtained from the ResNet output, where each element 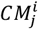 indicates the probability of contact between residues *i* and *j*.

#### 4.1.2. GAT-based binding pocket prediction model architecture

The GAT model serves as the foundational architecture for conducting residue-level classifications on protein graphs; this process specifically targets the prediction of binding pocket residues. After constructing the graph, where each node is characterized by an initial feature *P*^(0)^ and each edge is characterized by a feature 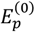, a GAT is employed to embed the constructed graphs into fixed-size latent representations, typically those residues involved in the binding pocket. The core of the model is built upon four GATConv layers. In each layer, the model aggregates information from its neighbouring nodes using an attention mechanism. Specifically, for the *l* − *th* layer, the feature update for node *i* is defined as shown below:

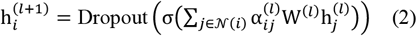

where 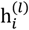 refers to the feature vector of node *i* at layer *l*, W^(*l*)^ is the learnable weight matrix at layer *l*, 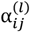 denotes the attention coefficient between node *i* and its neighbour *j* (with j ∈ N(i)), σ(·) is an ELU activation function that is applied after each convolution except that in the final layer, and dropout is used to randomly zero out a fraction of the activations to mitigate overfitting. The final convolutional layer, referred to as conv4, outputs logits for each node:

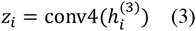

where *z*_*i*_ represents the unnormalized score produced for each class. These logits are subsequently transformed into probabilities via the Softmax function during the loss computation. Owing to the limited amount of training data (only 78 crystal PDBs; Supplementary file), we further incorporate data augmentation and consistency regularization. Specifically, Gaussian noise is added to the initial node embeddings to generate perturbed versions of the node features:

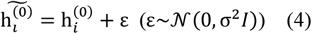

where *σ* controls the standard deviation of the noise.

To further enhance the robustness of the initial multilayer perceptron (MLP) classifier, we employ a consistency regularization-based semisupervised framework, as used by Huang et al[17b]. The datasets consisting of labeled and unlabeled A-domain structures are described in the Supplementary file. The overall training loss is defined as follows:

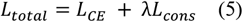

where *λ* is a hyperparameter that balances the contribution of the consistency regularization scheme relative to the cross-entropy loss. The detailed hyperparameter configurations are summarized in Table S3. Despite achieving 97.9% accuracy in pocket residue prediction using ABP-GAT, some errors remained. To correct residual misalignments, we applied a template-based post-processing step, in which a template file constructed by segmenting pocket sequences from the training set was used to calibrate the final pocket residue assignments.

#### 4.1.3 LoRA Fine-tuning of the ESM2 for A-Domain binding pocket characterization

To effectively characterize the 27-AA binding pockets of A-domains, we fine-tuned the ESM2 pretrained model (esm2_t33_650M_UR50D) using a masked language modeling task on a curated dataset of 191,303 ABP sequences (labeled and unlabeled), resulting in ABP-ESM2 (Supplementary File). The dataset was randomly split into training and validation sets with a 9:1 ratio. To enable multi-GPU distributed and mixed-precision training, we implemented the fine-tuning procedure in the PyTorch Lightning (v2.3.3) framework combined with the parameter-efficient tuning method LoRA [41]. We applied random masking (masking rate of 10%) to the ABP sequences for masked language modeling. The LoRA configuration was set as follows: rank (r = 16), scaling factor (alpha = 64), and dropout rate (0.1).

During training, we used the cross-entropy loss function computed only at masked positions for amino acid prediction and adopted the AdamW optimizer. The initial learning rate was dynamically adjusted based on validation performance using a plateau-based scheduler with a patience of 5 epochs and a decay factor of 0.5. Training was run for up to 500 epochs with early stopping based on validation loss, and the best-performing checkpoint was retained for subsequent analyses and applications. All training was conducted in data-parallel mode on NVIDIA A100 GPUs. The resulting ABP-ESM2 model is directly used to characterize the binding pocket sites of A-domains and to provide high-quality feature representations for downstream tasks.

### 4.2. SHAP-guided functional residue recognition in A-domain binding pockets

#### 4.2.1. A-domain binding pocket residue prediction and sequence analysis

The predicted ABP residues of 4545 A-domain were concatenated into continuous sequences and then aligned using MAFFT v7.310[42]. An ABP phylogenetic tree was constructed from the multiple sequence alignment with IQ-TREE v3.0.1[43] and visualized using iTOL[44]. To assess how well the phylogeny reflects substrate specificity, we cut the tree at a fixed height threshold (H = 2.0 substitutions) to obtain leaf clusters and compared these tree-derived clusters with known substrate labels using the adjusted Rand index (ARI)[28], where ARI = 1 indicates perfect agreement and ARI ≈ 0 is expected for random clustering. Within bacterial sequences, we additionally computed one-vs-rest per-label ARI for substrate labels represented by at least 2 sequences, and summarized results for labels with N ≥ 20, which were visualized as vertical bar plots in Python.

#### 4.2.2. TreeSHAP model construction process for functional and substrate-specific residue analyses

TreeSHAP[45], a variant of SHAP[22], leverages Shapley values from cooperative game theory to provide sample-level feature attributions for tree-based models (e.g., random forests and gradient-boosted trees), offering mathematically consistent and human-interpretable explanations, making it a gold-standard method in explainable artificial intelligence applications. For the TreeSHAP-based analysis, we first mitigated class imbalance in the 4,545 ABP sequences by random oversampling: for each substrate class with fewer than 10 samples, minority-class sequences were randomly duplicated until at least 10 sequences were available. This oversampled dataset was used only for the explainability analysis and did not affect training of the main predictive models. We then trained one-vs-rest random forest classifiers for each substrate class using 27-AA pocket features derived from ABP-ESM2, with 5-fold cross-validation and hyperparameter optimization via GridSearchCV (Table S3). For each fold, TreeSHAP was applied to the corresponding validation set to compute per-position feature contributions, and SHAP values from all folds were aggregated to obtain out-of-fold importance profiles for each substrate class. These TreeSHAP-based random forest models are used purely as post hoc interpretability tools and are not part of the primary prediction pipeline of DeepAden. The SHAP value for feature *i* of sample *x* belonging to class *c* is defined as follows:

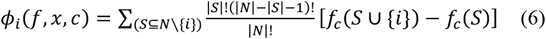

where *N* is the set of all features, *S* is a subset of the features excluding *i*, and *f*_*c*_ is the expected prediction of class *c* when only the features in set *S* are known. In the one-vs-rest setting, *f*_*c*_ denotes the output of the one-vs-rest random forest classifier corresponding to class *c*. We analysed the SHAP values across different classes to identify substrate-specific residues. For each site *i*, we calculated the mean absolute SHAP value across all classes:

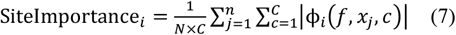

where *C* is the number of classes and *n* is the number of samples. For specific classes of interest, we compared their SHAP profiles with those of other classes to identify discriminative residues:

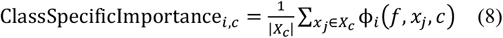

where *X*_*c*_ is the set of samples belonging to class *c*.

### 4.3. Contrastive learning for pocket–substrate binding prediction

#### 4.3.1. Pocket featurization (pocket encoder)

We generated the substrate features of the A-domains using ABP-ESM2. For a protein pocket with a length of *n*, the pocket encoder generated a pocket embedding 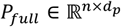 (*d*_*p*_=256), which was mean-pooled along the length of the pocket, resulting in a vector 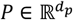. Notably, the pocket encoder was used exclusively to generate pocket features, with their weights remaining unchanged and not being updated during training.

#### 4.3.2. Substrate featurization (molecular encoder)

We generated the substrate features of A-domains using a pretrained CLM called MoLFormer[16b], encoding the SMILES string of a molecule as a fixed-dimensional embedding 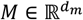, *d*_*m*_=768. Akin to the pocket encoder, MoLFormer served exclusively as a molecule feature generator, and its parameters were kept frozen throughout the training process.

#### 4.3.3. Projection and prediction

Given a pocket embedding 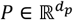 and a ligand embedding 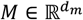, we projected them into a shared embedding space (*P*^*^, *M*^*^ ∈ ℝ^*d*^, *d*=128) using two separate two-layer MLPs:

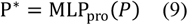

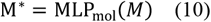

where 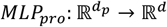 and 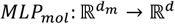 are two-layer perceptrons with rectified linear unit (ReLU) activation[46]. Given the latent embeddings *P*^*^, *M*^*^in the shared space, we employed cosine similarity as a metric to measure their binding affinity 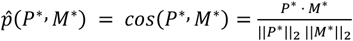. The intuition behind using cosine similarity is that pocket–ligand pairs with stronger binding affinities tend to exhibit higher cosine similarity values and are therefore closer in the embedding space.

#### 4.3.4. Training

The model was trained to predict the affinity between pockets and ligands, with the loss computation depending on the training dataset. To enhance the cross-modal alignment effect, we propose a contrastive learning framework that optimizes feature representations by simultaneously attracting genuine binding pairs in the embedding space while repelling nonbinding counterparts. Specifically, we define positive and negative sample pairs based on a batch of samples: given *N* protein pockets 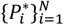 and their corresponding *N* molecules 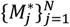, we combine them into *N*^2^ pocket–molecule pairs, where *i, j*∈[1, *N*]. A pair is considered positive if *i* = *j* or if the label of the pocket *y*_*i*_ matches the label of the molecule *y*_*j*_; it is considered negative if *i*≠*j* and *y*_*i*_≠*y*_*j*_. Based on this, we introduce two losses: Pocket-to-Mol loss and Mol-to-Pocket loss[47]. The former represents the likelihood that a given protein pocket 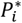 exhibits higher similarity to its positive molecules than to all negative molecules:

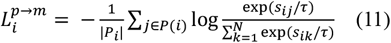

where *P*_*i*_ = {*j* | *y*_*i*_ = *y*_*j*_, *j* ∈ [1, *N*]} denotes the set of indices of all positive molecules for protein pocket 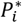, and 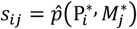 represents the normalized cosine similarity.

The latter defines the likelihood that a given molecule 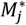 exhibits higher similarity to its positive pockets than to all negative pockets:

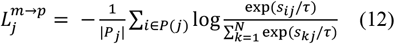

In both equations, τ denotes the temperature parameter controlling the softmax distribution. Combining the two losses, we obtain the final loss function for a batch:

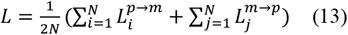

This bidirectional objective, calibrated by temperature scaling, directly optimizes the discriminative boundaries between paired instances. Such a formulation makes the loss particularly suitable for supervised contrastive learning in tasks requiring precise interaction modeling. The detailed implementation specifications and parameter configurations are provided in Table S3.

#### 4.3.5. Probability calibration of model predictions

In the inference stage, DeepAden applies a kernel density estimation (KDE)–based calibration method to convert cosine similarity scores into statistically meaningful confidence probabilities[23]. Using the augmented training data, it computes the similarity between each protein pocket and all molecules to obtain the empirical similarity distributions for positive and negative pairs. These two distributions are modeled separately in a non-parametric manner using Gaussian-kernel KDE, where the bandwidth is set by Scott’s rule and further scaled by a smoothing factor (α = 2) to yield a smoother and more robust estimate. During inference, DeepAden calculates the cosine similarity between a query protein pocket and all molecular embeddings in the library, ranks the candidate molecules in descending order of similarity, and then uses the calibrated positive and negative similarity densities within a Bayesian framework to map each similarity score to a posterior probability of being a positive sample. In this way, raw similarity scores are converted into statistically interpretable confidence scores, providing reliable quantification of prediction uncertainty.

### 4.4. Experimental validation

#### 4.4.1. Actinomycete material, culture conditions, and DNA preparation

*Streptomyces hygroscopicus* OsiSh-2 (GenBank accession JBPQZZ000000000)[32] was originally isolated from the sheath of healthy rice (Oryza sativa cv. Gumei 4) collected in Liuyang, Hunan Province, China. For the primary cultivation task, the strain was grown on Gauze agar plates at 28°C for 5 days. A seed culture was subsequently prepared by inoculating the strain into liquid Gauze medium, followed by incubation at 28°C with agitation at 180 rpm for 5 days. The cultured broth was then transferred to Gauze plate medium for solid-state fermentation at 28°C for 7 days. High-quality genomic DNA of *Streptomyces hygroscopicus* OsiSh-2 was extracted using a standardized protocol referred to Huang et al[17b].

#### 4.4.2. LC-MS/MS analysis and molecular networking

Crude extracts obtained from different fermentation methods were processed and analysed by LC-MS/MS. Chromatographic separation was performed using an Agilent G6500 UHPLC system coupled to a quadrupole time-of-flight (Q-ToF) mass spectrometer operating in the positive ion mode. The samples (5 μL injection volume) were separated on a Phenomenex Kinetex C18 column (1.7 μm particle size,100 × 2.1 mm, 100 Å) with a flow rate of 0.3 mL/min. The mobile phase consisted of (A) 0.1% formic acid in water and (B) 0.1% formic acid in acetonitrile (ACN), with the following gradient program: 20% B (0–2 min), a linear increase to 100% B (2–14 min) and holding 100% B (14–16 min), followed by re-equilibration at 20% B for 2.8 min (total run time: 18.8 min). The spray chamber conditions were as follows: nebulizer: 5 L/min; drying gas: 200; sheath gas temperature: 350°C; sheath gas flow: 11 L/min; and drying gas on: 5 L/min.

Raw LC-MS data derived from the OsiSh-2 fermentation extracts were processed using the Global Natural Product Social Molecular Network (GNPS) platform[48]. Molecular networks were constructed and visualized using Cytoscape v3.9.1[49] with the following parameters: precursor and fragment ion mass tolerances of 0.02 Da, a minimum cosine score of 0.6, at least two matched fragment ions, and a minimum cluster size of 1. Unidentified metabolites were further annotated by querying the NPAtlas[50], Reaxys, and SciFinder databases.

## Acknowledgements

This work was supported by the National Natural Science Foundation of China (32170079 to Z.Q., 32200035 to H.Z., and 32400235 to J.H.), the Natural Science Foundation of Guangdong (2024A1515012593 to Z.Q. and 2023A1515110175 to J.H.), Guangdong Talent Scheme (2021QN020100 to Z.Q.).

## Conflict of Interest

The authors declare no competing financial interests.

## Author Contributions

Zhiwei Qin and Heqian Zhang designed and supervised the research. Jiaquan Huang, Liangjun Ge and Yaxin Wu performed the bioinformatic and established the algorithm. Song Meng, Qiandi Gao and Pan Li participated the bioinformatic analysis. Jun Wu performed experiment validation. All authors analysed and discussed the data. Zhiwei Qin, Heqian Zhang, Jiaquan Huang and Liangjun Ge wrote the manuscript and all authors edited.

## Data Availability Statement

The data and materials supporting the findings reported in this study are available in the supplementary material of this article. The code is available at GitHub repository: https://github.com/Qinlab502/DeepAden. We also developed a website that allows users from any computer background to use it, which can be accessed at https://deepnp.site/. The Web server usage and parameter settings was supplemented in in the supplementary material.

## Notes

### Competing Interest Statement

The authors have declared no competing interest.

### Summary of Updates

This revised version incorporates changes made in response to the peer reviewers comments. All figures and figure legends have been updated. In addition, the manuscript has been extensively revised to improve clarity, readability, and accessibility for a broader readership.

