## Supplementary material for "DeepAden: An explainable machine learning method for predicting the substrate specificity of nonribosomal peptide synthetases": materials and methods

### 1. Data collection and processing

#### 1.1 A-domain sequence data with known labels

We collected a total of 4,586 A-domain sequences encompassing 247 distinct substrate labels from three sources: the MIBiG database (Version 3.1)[1] containing experimentally validated or structurally supported data points (1,627 sequences), the PARAS dataset[2] (3,257 sequences), and the NRPStransformer dataset[3] (4,430 sequences) **(Dataset S1)**. To prepare the data for the contrastive learning-based substrate prediction model training, we implemented a rigorous data processing pipeline: (1) Sequence redundancy removal was performed using CD-HIT (version 4.8.1)[4] with a 95% sequence identity threshold, yielding 4,627 non-redundant A-domain sequences across the three datasets. (2) Sequences lacking substrate labels or containing "branch" annotations were excluded. (3) Synonymous labels were standardized, and erroneous substrate annotations were corrected through literature curation. (4) To ensure compatibility with the molecular language model MolFormer[5], which does not distinguish molecular stereoisomers, all substrate SMILES representations were canonicalized using RDKit (http://www.rdkit.org). (5) Since many data points possessed multiple substrate labels potentially arising from post-A-domain modifications by tailoring enzymes (e.g., methyltransferases, hydroxylases, halogenases) within biosynthetic gene clusters (BGCs), we excluded A-domain sequences with multi-substrate labels likely attributable to such modifications. The remaining multi-substrate entries were merged with single-substrate data, ultimately yielding 4,545 A-domain sequences spanning 223 substrate labels for training the substrate prediction model **(Dataset S2、S3)**.

#### 1.2 A-domain sequence data with unknown labels

To comprehensively identify bacterial-derived NRPS A-domain sequences for subsequent fine-tuning of the ABP-ESM model, we implemented a systematic sequence collection strategy. First, all bacterial genomes were retrieved from the Genome Taxonomy Database (GTDB Release 220)[6], and BGCs were predicted using antiSMASH (v7.0)[7]. Coding sequences (CDSs) encoding nonribosomal peptides (NRPs) were then extracted, generating a large pool of putative NRP gene cluster sequences. A-domains within these sequences were isolated through sequence alignment against known A-domain described above with Diamond (v2.1.11)[8], yielding 82,748 successfully aligned A-domain sequences. In parallel, we retrieved 152,502 A-domain sequences from the UniRef database (release-2024_06). Following merging and deduplication of these two datasets, 233,914 unique sequences were obtained. These sequences were subsequently validated through HMM-based matching against NRPS A-domain profiles (Pfam IDs: PF00501.29 and PF13193.7) using HMMER (v3.3.2). After stringent filtering and deduplication, we ultimately obtained 186,758 metagenomic A-domain sequences with an average length of 495 amino acids.

#### 1.3 A-domain structure data

A comprehensive dataset of 78 high-quality three-dimensional A-domain structures was curated from the Protein Data Bank (PDB) for model development. Initially, 10 A-domain structures with substrate co-crystallization were selected from the PDB database as the core training set. To enhance dataset diversity and model generalization capability, we expanded the initial collection by retrieving homologous structures based on stringent similarity criteria: sequence identity >30% and root-mean-square deviation (RMSD) <3 Å. Following quality assessment and redundancy removal, the final curated dataset comprised 78 non-redundant, high-quality three-dimensional A-domain structures **(Dataset S4)**, which were subsequently utilized to train the residue contact prediction model and the graph attention network (GAT)-based binding pocket prediction model. For structures with co-crystallized substrates, active pockets were defined as residues within 6 Å of the substrate, spanning four discontinuous regions that collectively comprised 27 amino acids. For the remaining structures, the corresponding active pocket residues were determined through structural alignment using the align command in PyMOL (version 3.1.0). Finally, the labelled active pocket data derived from 78 experimentally validated protein structures were used for supervised training, and 1,474 AlphaFold3-predicted protein structures were used for semisupervised learning.

### 2. Construction of ResNet-based residue contact prediction model

#### 2.1 Model Architecture

We developed a deep learning framework for predicting binary residue-residue contacts from protein sequences, utilizing evolutionary-scale information from protein language models[9]. Our architecture comprises two main components: (1) a pretrained protein language model serving as a feature extractor, and (2) a specialized residual projection histogram network for contact map prediction.

##### 2.1.1 Feature extraction with evolutionary scale modeling (ESM)

We employed ESM2 (Evolutionary Scale Modeling 2) as our backbone feature extractor. Specifically, we utilized the esm2_t33_650M_UR50D variant containing 33 transformer layers with 650 million parameters. For each input protein sequence $S$ of length $L$, we obtained:

$$\mathrm{E}, \mathrm{A} = \mathrm{ESM}(S)$$

where $\mathrm{E} \in \mathbb{R}^{L\times d}$ represents the residue-wise embeddings ($d$=1280), and $\mathrm{A} \in \mathbb{R}^{33\times 20\times L\times L}$ denotes the multi-head attention maps across all transformer layers and attention heads. These attention maps capture pairwise dependencies between residues at different abstraction levels.

##### 2.1.2 Residual projection distogram network

To transform attention maps into contact predictions, we designed a residual projection histogram network. The attention maps from all layers and heads (A) are flattened along the layer dimension, resulting in $\mathrm{A}' \in \mathbb{R}^{660\times L\times L}$. A series of six residual blocks progressively reduce feature dimensions: 660→512→256→128→64→32→16 channels. Each residual block contains two convolutional layers with batch normalization and ReLU activations, with skip connections to mitigate vanishing gradients. A final 1×11×1 convolution produces logits $\mathrm{O} \in \mathbb{R}^{L\times L}$, which are symmetrized to ensure consistency:

$$\mathrm{O}_{\mathrm{sym}} = \frac{1}{2}(\mathrm{O} + \mathrm{O}^{\mathsf{T}})$$

The output is passed through a sigmoid function to obtain contact probabilities $\mathrm{P} \in [0, 1]^{L\times L}$.

##### 2.1.3 Objective function

We formulated contact prediction as a binary classification problem. Given ground truth contact matrix $\mathrm{Y} \in \{0, 1\}^{L\times L}$ (where contacts are defined as residue pairs with $C_\beta$-$C_\beta$ distance < 8Å), we minimized the binary cross-entropy loss (BCE loss).

#### 2.2 Training Procedure

#### 2.2.1 Dataset Preparation

We curated a dataset of 78 A-domain sequences with experimentally determined structures from the Protein Data Bank. Contact maps were generated using $C_\beta$ atom distances with an 8Å cutoff threshold. The dataset was randomly partitioned into training (80%) and validation (20%) sets with stratification to ensure balanced representation.

#### 2.2.2 Optimization

The model was trained using AdamW optimizer[10] with initial learning rate 1e-3, weight decay 1e-2, and batch size 1. We employed a ReduceLROnPlateau scheduler that reduced the learning rate by a factor of 0.8 when the validation Matthews correlation coefficient (MCC) plateaued for 15 epochs, with a minimum learning rate of 1e-6.

#### 2.2.3 Evaluation Metrics

Model performance was assessed using multiple metrics calculated per-protein and averaged:

$$\text{Accuracy (ACC)} = \frac{TP \times TN - FP \times FN}{TP + TN + FP + FN}$$

$$\text{Precision (PRE)} = \frac{TP}{TP + FP}$$

$$\text{Recall (REC)} = \frac{TP}{TP + FN}$$

$$\text{F1-score} = 2 \times \frac{\text{PRE} \times \text{REC}}{\text{PRE} + \text{REC}}$$

$$\text{Matthews Correlation Coefficient (MCC)} = \frac{TP \times TN - FP \times FN}{\sqrt{(TP + FP)(TP + FN)(TN + FP)(TN + FN)}}$$

We implemented a dynamic threshold optimization procedure, evaluating thresholds from 0 to 1 in 0.05 increments to maximize MCC on the validation set. The best-performing model checkpoint was selected based on validation MCC.

#### 2.3 Implementation Details

All experiments were conducted using PyTorch 2.1.0 with CUDA 12.0 acceleration on a single NVIDIA A100 GPU. The model parameters were initialized from pretrained ESM2 weights, with the convolutional layers initialized using Kaiming initialization[11]. Code is available at https://github.com/Qinlab502/DeepAden/blob/main/train/CM_trainning.py.

### 3. Usage of DeepAden web server

The DeepAden web server accepts input in the form of amino acid sequences in FASTA format, genomic files in FASTA format, and GBK files. Users can adjust the top-k parameter to select the top k predictions for each sequence (default k=3). In addition to our predefined database of 223 substrate molecules, users may upload their own custom molecular database as a CSV file with columns labeled "label" and "smiles" (canonical SMILES format is recommended). To enhance the user experience, after model evaluation, we retrained DeepAden using all available data including 4,545 train data and 201 A-domain sequences from the evaluation dataset, resulting in the model weight file "all.weight." To ensure the reproducibility of the results in this work, we also provide the model weights used during evaluation, stored as "benchmark.weight."

The prediction results display the top k predicted substrates along with their corresponding confidence scores (ranging from 0 to 1), with each score being independent of the others. We observed that when the top-1 prediction score is low (<0.5), the true substrate often appears among the top three predictions with a relatively low score. For cases involving multiple substrates, the model typically predicts the primary substrate with high confidence (1.0), while secondary substrates may appear in the top three predictions with lower scores. We offer multiple output formats for downloading results, including CSV and JSON, to facilitate the use of predictions in downstream tasks.

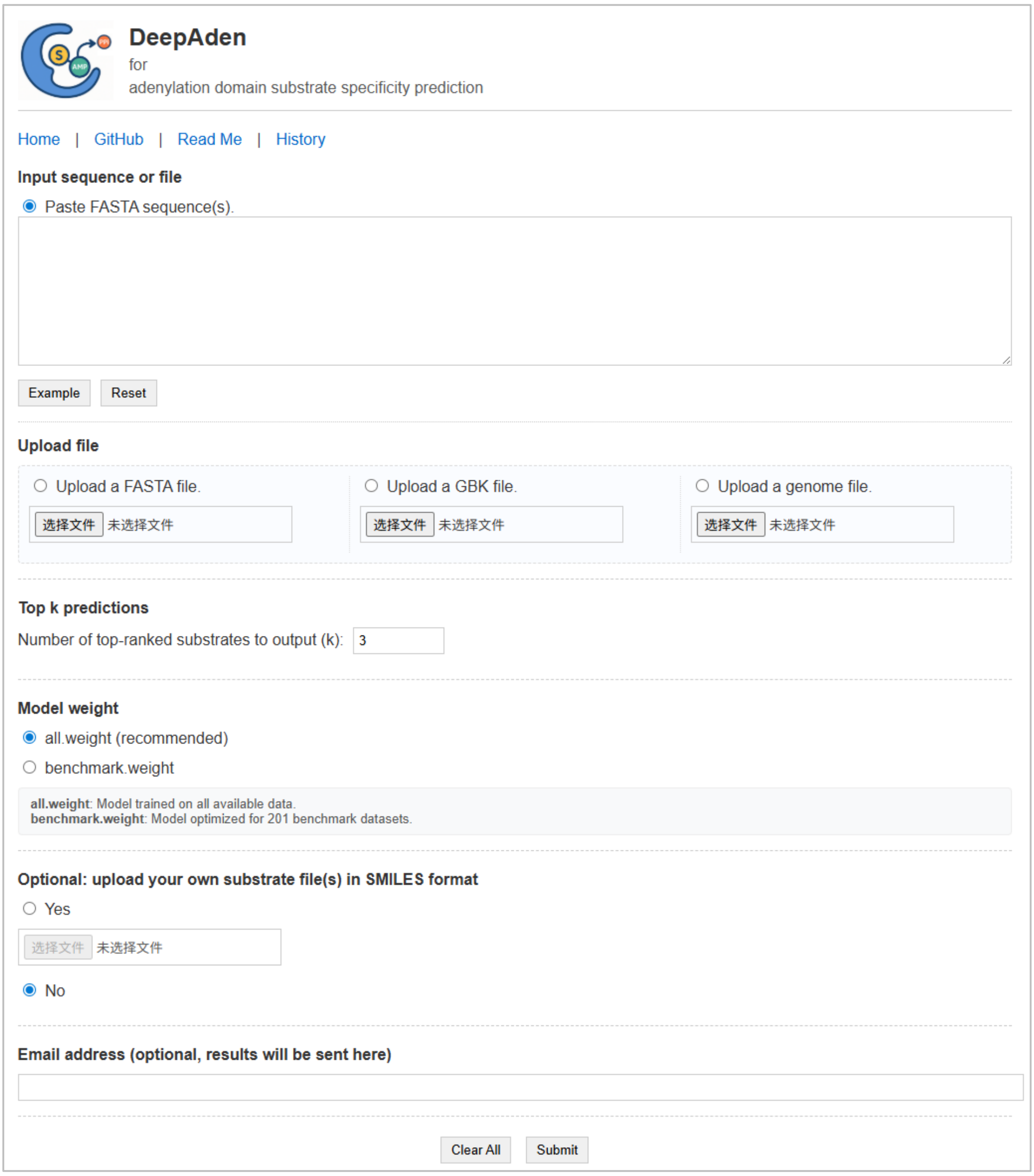


**Figure-**User interface of DeepAden web server

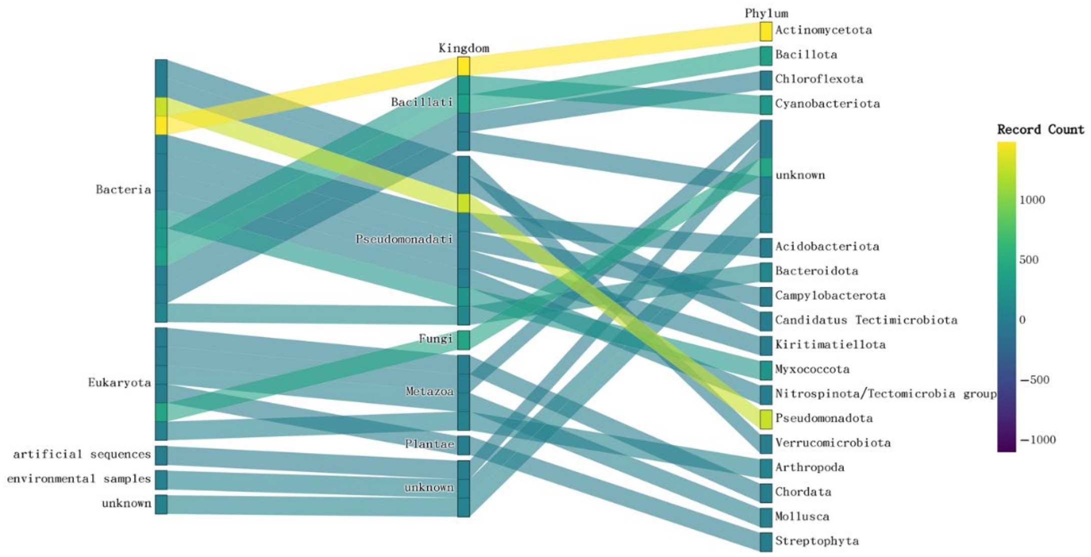


**Figure S1.** The 4545 A-domain substrates information and organism sources of A-domain data used in our research.

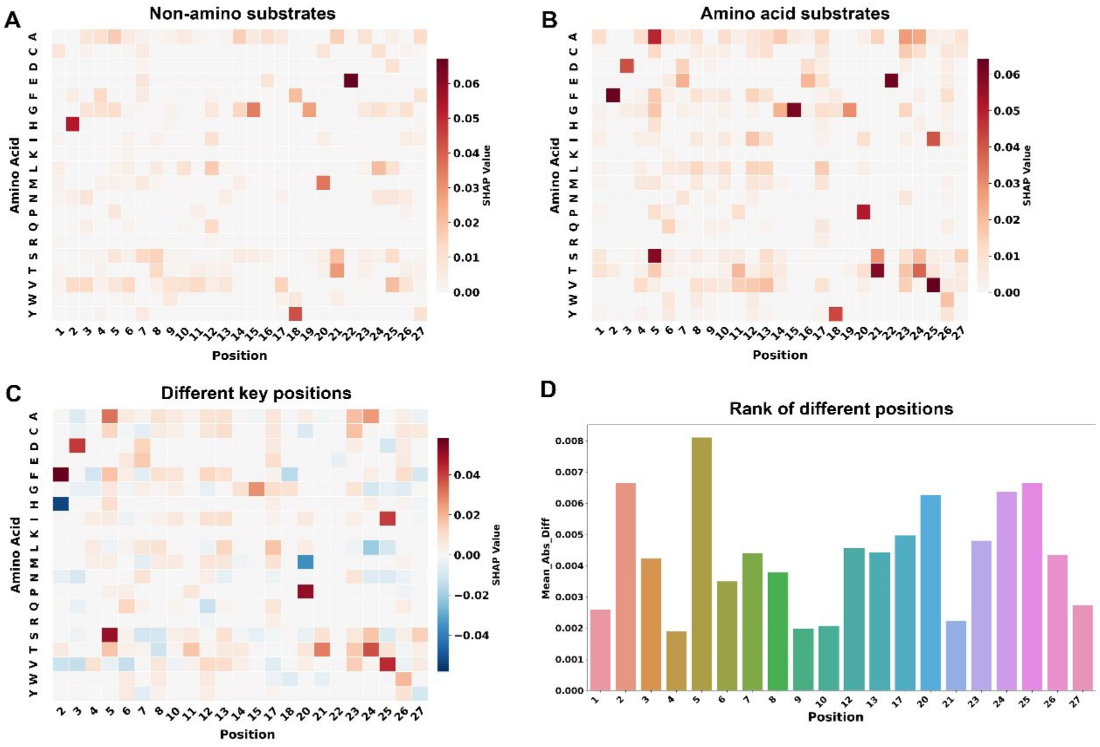


**Figure S2.** Heatmaps showing positive SHAP values that highlight amino acid contributions at each residue position for (**A**) non-amino ABPs (Dhb, PABA, Sal, Kiv, Cia, Pa, Ana, Ppa, 4-APCA, Qca, Box) and (**B**) amino acid substrate ABPs (Cys, Ser, Val, Gly, Pro, Ala, Thr, His, Asn, Gln, Tyr) within their respective classes. (**C**) Heatmap displaying the differences in key residue contributions between non-amino ABPs and amino acid substrate ABPs. (**D**) Bar plot ranking the SHAP value differences between non-amino ABPs and amino acid substrate ABPs.

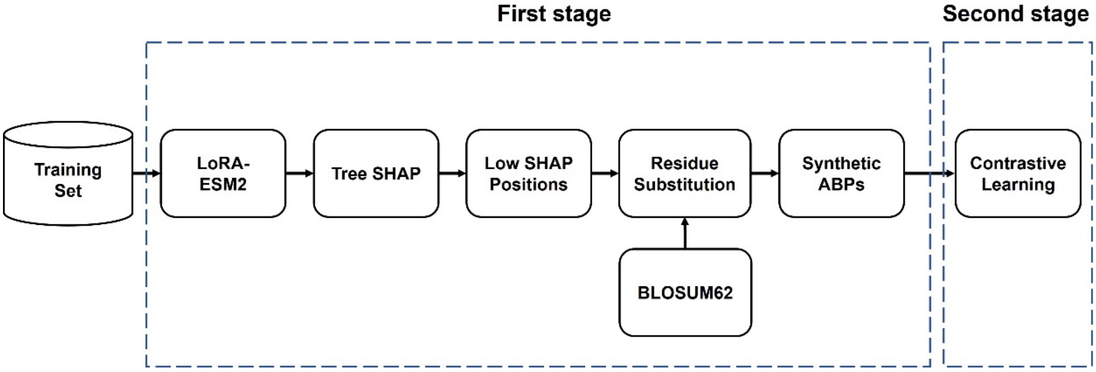


**Figure S3.** Flowchart of SHAP-guided data augmentation for contrastive learning

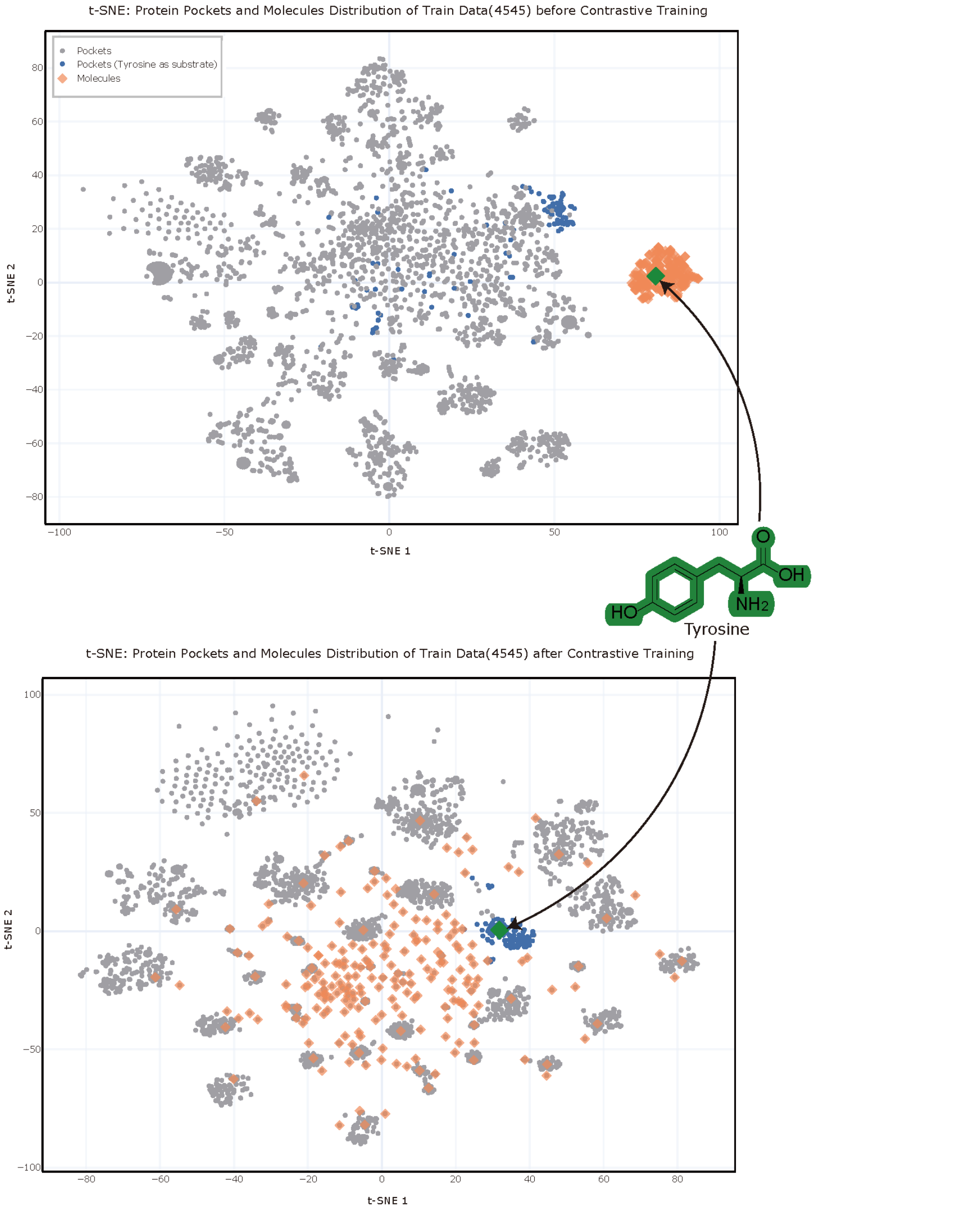


**Figrue S4.** t-SNE visualization of 4545 A-doamin pocket sequences (dots) and their corresponding 223 substrate molecules (diamonds). Before contrastive training, the substrate molecule Tyrosine (green diamond) is distant from its corresponding pockets (blue dots) in the vector space. After contrastive training, Tyrosine and its corresponding pockets become closely clustered.

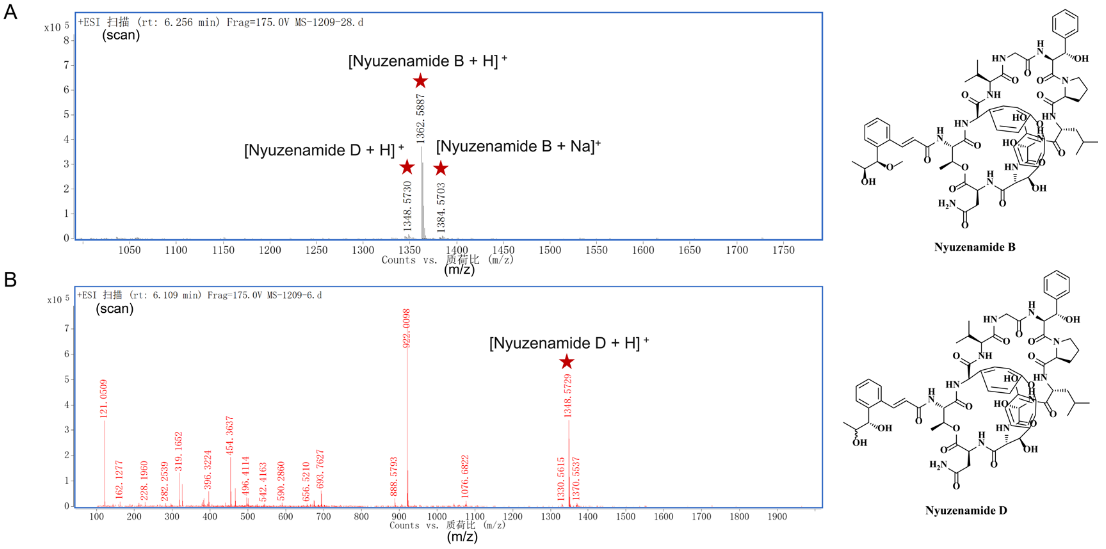


**Figure S5.** LC–MS and MS/MS analysis of nyuzenamides B and D from *Streptomyces hygroscopicus* OsiSh-2. **(A)** Extracted mass spectra showing the main ions assigned to nyuzenamides B and D, including [nyuzenamides B + H]^+, [nyuzenamides D + H]^+ and the sodium adduct [nyuzenamides B + Na]^+. **(B)** MS/MS spectrum of the nyuzenamide D precursor, which is dominated by precursor and neutral-loss ions with only limited backbone cleavage, in line with the rigid, bicyclic scaffold and stable ester linkages. The fragmentation pattern closely matches the GNPS reference spectrum of NMR-validated nyuzenamide D. In addition, DeepAden predictions for the A-domain substrate specificities in BGC 4.24 show a high degree of agreement with the reported nyuzenamide peptide backbone, further supporting the assignment of these metabolites as nyuzenamides produced by strain OsiSh-2.

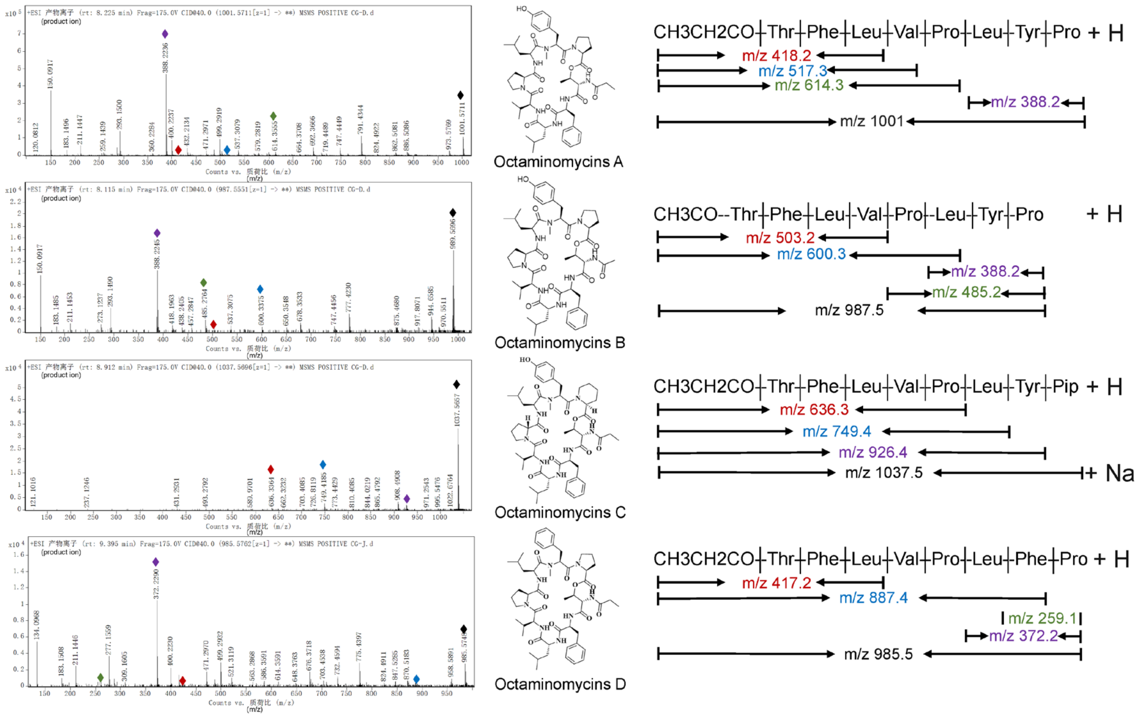


**Figure S6.** MS/MS spectra and diagnostic fragment ions used to establish the peptide backbones of octaminomycins A – D. (Left) Representative positive-mode MS/MS spectra of octaminomycins A – D, with the major sequence-informative ions highlighted. (Right) Summary of key fragment ions (color-coded by compound) whose m/z values correspond to consecutive N- or C-terminal peptide segments. These ions collectively confirm the proposed backbones and distinguish the four octaminomycin analogs from one another.

Select genomic region:
Overview 4.1 4.2 4.3 4.4 4.5 4.6 4.7 4.8 4.9 4.10 4.11 4.12 4.13 4.14 4.15 4.16 4.17 4.18 4.19 4.20 4.21 4.22 4.23 4.24 4.25 4.26 4.27 4.28 4.29 4.30 4.31 4.32 4.33 4.34 4.35 4.36 4.37 4.38
4.41 4.42 5.1

**Identified secondary metabolite regions using strictness 'relaxed'**

| Region | Type | From | To | Similarity Confidence | Most similar known cluster | |
|---|---|---|---|---|---|---|
| Region 4.1 | T1PKS | 1 | 45,358 | Low | hygromycin A | saccharide |
| Region 4.2 | NRP-metallophore, NRPS, NRPS-like, betalactone, T1PKS, T3PKS | 82,996 | 244,645 | Medium | totopotensamide A/totopotensamide B | NRPS:Type I+PKS |
| Region 4.3 | terpene | 441,914 | 463,050 | Low | actinospectacin | saccharide |
| Region 4.4 | terpene | 690,904 | 716,693 | High | isorenieratene | terpene |
| Region 4.5 | NRPS-like | 727,861 | 769,813 | | | |
| Region 4.6 | T1PKS | 839,776 | 979,681 | Medium | azalomycin F3a | PKS |
| Region 4.7 | terpene | 1,109,294 | 1,130,529 | | | |
| Region 4.8 | T1PKS, NRPS | 1,142,250 | 1,351,916 | High | nigericin | PKS:Type I |
| Region 4.9 | hserlactone | 1,580,122 | 1,600,883 | | | |
| Region 4.10 | butyrolactone | 1,605,489 | 1,616,424 | | | |
| Region 4.11 | NAPAA | 2,057,465 | 2,091,601 | Low | γ-poly-L-2,4-diaminobutyric acid | NRPS:Type I |
| Region 4.12 | lanthipeptide-class-i | 2,095,335 | 2,119,776 | Low | steffimycin D | PKS:Type II+saccharide:hybrid/tailoring |
| Region 4.13 | terpene | 2,293,433 | 2,319,961 | High | hopene | terpene |
| Region 4.14 | T2PKS | 2,473,962 | 2,546,477 | High | spore pigment | PKS |
| Region 4.15 | PKS-like, transAT-PKS, NRPS, NRPS-like, redox-cofactor | 2,599,787 | 2,704,451 | Medium | Largimycin A1/Largimycin A2/Largimycin A3 | NRPS:Type I+PKS:Type I |
| Region 4.16 | RiPP-like | 2,848,695 | 2,859,600 | | | |
| Region 4.17 | ectoine | 2,943,246 | 2,953,644 | High | ectoine | other:other |
| Region 4.18 | NI-siderophore | 3,033,842 | 3,063,638 | | | |
| Region 4.19 | NRPS-like | 3,603,149 | 3,645,995 | High | echoside A/echoside B/echoside C/echoside D/echoside E | NRPS:Type I |
| Region 4.20 | NI-siderophore, T2PKS | 4,159,471 | 4,237,041 | High | enterocin | PKS:Type II |
| Region 4.21 | T1PKS, NRPS | 4,445,308 | 4,568,264 | High | alchivemycin A/alchivemycin B | PKS+NRPS:Type I |
| Region 4.22 | terpene | 5,300,807 | 5,323,260 | High | geosmin | terpene:Sesquiterpene |
| Region 4.23 | arylpolyene | 5,561,458 | 5,602,591 | Low | skyllamycin D/skyllamycin E | NRPS:Type I |
| Region 4.24 | NRPS | 5,605,917 | 5,688,015 | Low | skyllamycin A/skyllamycin B | NRPS:Type I+PKS |
| Region 4.25 | indole | 6,013,493 | 6,034,638 | Medium | 5-isoprenylindole-3-carboxylate β-D-glycosyl ester | other:other |
| Region 4.26 | terpene, NRPS | 6,224,379 | 6,285,460 | Medium | ochronotic pigment | other:other |
| Region 4.27 | HR-T2PKS | 6,330,995 | 6,381,478 | Low | (2E,4E,6E,8E)-N-(2-hydroxy-5-oxocyclopent-1-en-1-yl)-9-(o-tolyl)nona-2,4,6,8-tetraenamide/(E)-N-(2-hydroxy-5-oxocyclopent-1-en-1-yl)-5-(o-tolyl)pent-4-enamide | PKS+NRPS:Type I |
| Region 4.28 | RiPP-like | 6,388,978 | 6,401,500 | | | |
| Region 4.29 | T1PKS, terpene-precursor | 6,541,649 | 6,741,169 | Medium | desulfoclethramycin/clethramycin | other:other |
| Region 4.30 | ectoine | 7,712,953 | 7,723,357 | High | ectoine | other:other |
| Region 4.31 | NI-siderophore | 7,878,234 | 7,910,914 | Low | peucechelin | NRPS:Type I |
| Region 4.32 | terpene | 7,977,135 | 7,999,063 | | | |
| Region 4.33 | NRPS-like, deazapurine, NRPS | 8,115,635 | 8,163,925 | Low | toyocamycin | other:other |
| Region 4.34 | T1PKS, amglyccycl | 8,243,095 | 8,298,103 | Low | validamycin A | other:cyclitol+saccharide |
| Region 4.35 | T1PKS | 8,432,128 | 8,508,784 | High | butyrolactol A | PKS |
| Region 4.36 | T1PKS, NRPS-like | 8,609,704 | 8,689,423 | Medium | geldanamycin | PKS |
| Region 4.37 | terpene | 8,740,149 | 8,762,148 | High | cyclooctatin | terpene:Diterpene |
| Region 4.38 | terpene | 8,778,308 | 8,799,486 | High | pristinol | terpene |
| Region 4.39 | terpene-precursor, NRPS-like | 9,030,811 | 9,085,953 | | | |
| Region 4.40 | RiPP-like | 9,171,331 | 9,181,549 | | | |
| Region 4.41 | NRPS, lanthipeptide-class-i | 9,186,165 | 9,275,988 | | | |
| Region 4.42 | hydrogen-cyanide | 9,450,735 | 9,463,563 | Low | aborycin | ribosomal:RiPP |
| Region 5.1 | 2dos | 181,354 | 209,308 | Medium | hygromycin A | saccharide |

**Figure S7.** Analysis with antiSMASH 7.0[7] identified 12 NRPS biosynthetic gene clusters (indicated by red boxes) in *Streptomyces hygroscopicus* OsiSh-2.

**Table S1**. Construction of the 49 amino acids (49-AA) A-domain binding pocket within 8Å by comparing 10 co-crystal complexes of A-domains.

| PDB ID | Region 1 | Region 2 | Region 3 | Region 4 | Region 5 | substrate | Organism |
|---|---|---|---|---|---|---|---|
| **1amu** | ---------L--<br>FF----- | A---FDASVWE--M | -TL-- | -ITAGS | -NAYGPTETTICATT | phe | *Brevibacillus brevis* |
| **1md9** | --R----Y--S--<br>-S---- | L---HNYPLSSPG- | -ALV- | -QVGGA | -QVFGMAEGLVNYTR | dhb | *Bacillus subtilis* |
| **2vsq** | ---------L---<br>------ | SN-AFDAFTFDFYA | -FATT | -LFGGE | -NCYGPT--TVFATA | leu | *Bacillus subtilis* |
| **4d56** | ---------F---<br>-----Y | V---FDVA-EE--- | WSLPT | VIIGGE | INCYGPTEGTIAVSL | tyr | *Planktothrix agardhii* |
| **4zxi** | -------------<br>------ | ----FDI------- | ----- | ---GGE | --VYGPTETTVWSSA | gly | *Acinetobacter baumannii* AB307-0294 |
| **5n9x** | -------------<br>------ | H---FDFS-W---- | -NQT- | -VFGGE | -NMYGITETTVHATF | thr | *Streptomyces* sp. |
| **7xbu** | ---------L--<br>W------ | A---FDPS-QQ--- | -DLVT | -IIGGE | NTIYGPTEAAVNAT- | cap | *Saccharothrix mutabilis subsp. capreolus* |
| **5wm2** | I-R----Y--N--<br>-S---- | L---HNFALACP-- | -AVV- | -QVGGS | -QVFGMAEGLLNY-- | sal | *Streptomyces gandocaensis* |
| **3vnr** | -------------<br>------ | H---FDFSVWE--- | -NQTP | -IFGGE | -NGYGITETTVFTTF | aba | *Streptomyces sp.* |
| **8gic** | ---------L---<br>--W--- | AP--FDASLFE--- | -HLTA | -LTGGD | RHLYGPTETTLCATW | hpg | *Actinoplanes teichomyceticus* |
| **49 AA** | X-X----X-XX-<br>XXXX--X | XX-XXXXXXXXXXX | XXXXX | XXXXXX | XXXXXXXXXXXXXXX | - | - |

**Table S2.** Comparison of amino acid residue positions in the 10-AA and 34-AA codes with the A-domain binding pocket (ABP) residues. All residue numbering is based on GrsA_Phe (PDB: 1amu). Residues highlighted in blue shadow represent informative positions identified by Terlouw et al[2], while residues highlighted in yellow shadow represent contributory positions identified in this study.

| 34-AA[14] | 27-AA | 10-AA[15] |
|---|---|---|
| 210 | | |
| 213 | | |
| 214 | | |
| 230 | **230** | |
| 234 | **234** | |
| 235 | **235** | 235 |
| **236** | **236** | 236 |
| 237 | 237 | |
| 238 | | |
| **239** | 239 | 239 |
| | 240 | |
| 243 | | |
| | 277 | |
| **278** | **278** | 278 |
| 279 | 279 | |
| | 280 | |
| 299 | 299 | 299 |
| 300 | 300 | |
| **301** | 301 | 301 |
| 302 | 302 | |
| 303 | 303 | |
| 320 | | |
| 321 | | |
| **322** | 322 | 322 |
| 323 | 323 | |
| 324 | 324 | |
| 325 | **325** | |
| 326 | 326 | |
| 327 | 327 | |
| 328 | 328 | |
| 329 | **329** | |
| 330 | 330 | 330 |
| 331 | 331 | 331 |
| 332 | **332** | |
| 333 | | |
| 334 | | |
| | | 517 |

**Table S3.** Training parameters for each module in DeepAden

| Model | Hyperparameters |
|---|---|
| Random forest-based A-domain binding pocket | n_estimators: 200<br>max_depth: None<br>min_samples_split: 2<br>min_samples_leaf: 1<br>bootstrap: False |
| Graph attention neural network-based A-domain binding pocket prediction model (ABP-GAT) | GAT model structure |
| | hidden_channels: 64<br>out_channels: 3<br>heads: 32<br>heads_intermediate: 16<br>heads_final: 8<br>dropratio: 0.5 |
| | Training |
| | learning rate: 1e-3<br>epoch: 700<br>early_stop_epochs: 200<br>batch_size: 8 |
| | Semi-supervised and data augmentation |
| | dropnode_rate: 0.4<br>dropedge_rate: 0.4<br>tem: 0.5<br>lam_initial: 0.1<br>lam_final: 0.5<br>lam_rampup_epochs: 300<br>order: 1 |
| Multimodal contrastive learning-based pocket-substrate binding prediction model | epochs: 100<br>learning_rate: 1e-4<br>batch_size: 512<br>optimizer: adamw<br>weight_decay: 1e-4<br>temperature(τ): 0.1<br>early_stopping_patience: 10<br>early_stopping_delta: 1e-4 |

### References


[1] a) Mitja M. Zdouc, K. Blin, Nico L. L. Louwen, J. Navarro, C. Loureiro, Chantal D. Bader, Constance B. Bailey, L. Barra, Thomas J. Booth, Kenan A. J. Bozhüyük, José D. D. Cediel-Becerra, Z. Charlop-Powers, Marc G. Chevrette, Y. H. Chooi, Paul M. D'Agostino, T. de Rond, E. Del Pup, Katherine R. Duncan, W. Gu, N. Hanif, Eric J. N. Helfrich, M. Jenner, Y. Katsuyama, A. Korenskaia, D. Krug, V. Libis, George A. Lund, S. Mantri, Kalindi D. Morgan, C. Owen, C.-S. Phan, B. Philmus, Zachary L. Reitz, Serina L. Robinson, K. S. Singh, R. Teufel, Y. Tong, F. Tugizimana, D. Ulanova, Jaclyn M. Winter, C. Aguilar, Daniel Y. Akiyama, Suhad A. A. Al-Salihi, M. Alanjary, F. Alberti, G. Aleti, Shumukh A. Alharthi, Mariela Y. A. Rojo, Amr A. Arishi, Hannah E. Augustijn, Nicole E. Avalon, J A. Avelar-Rivas, Kyle K. Axt, Hellen B. Barbieri, Julio Cesar J. Barbosa, L. G. Barboza Segato, Susanna E. Barrett, M. Baunach, C. Beemelmanns, D. Beqaj, T. Berger, J. Bernaldo-Agüero, Sandra M. Bettenbühl, Vincent A. Bielinski, F. Biermann, Ricardo M. Borges, R. Borriss, M. Breitenbach, Kevin M. Bretscher, Michael W. Brigham, L. Buedenbender, Brodie W. Bulcock, C. Cano-Prieto, J. Capela, Victor J. Carrion, Riley S. Carter, R. Castelo-Branco, G. Castro-Falcón, Fernanda O. Chagas, E. Charria-Girón, A. A. Chaudhri, V. Chaudhry, H. Choi, Y. Choi, R. Choupannejad, J. Chromy, Melinda S. C. Donahey, J. Collemare, Jack A. Connolly, Kaitlin E. Creamer, M. Crüsemann, Andres A. Cruz, A. Cumsille, J.-F. Dallery, Luis C. Damas-Ramos, T. Damiani, M. de Kruijff, B. D. Martín, G. D. Sala, J. Dillen, Drew T. Doering, Shravan R. Dommaraju, S. Durusu, S. Egbert, M. Ellerhorst, B. Faussurier, A. Fetter, M. Feuermann, David P. Fewer, J. Foldi, A. Frediansyah, Erin A. Garza, A. Gavriilidou, A. Gentile, J. Gerke, H. Gerstmans, J. P. Gomez-Escribano, Luz A. González-Salazar, Natalie E. Grayson, C. Greco, Juan E. G. Gomez, S. Guerra, S. G. Flores, A. Gurevich, K. Gutiérrez-García, L. Hart, K. Haslinger, B. He, T. Hebra, Jethro L. Hemmann, H. Hindra, L. Höing, Darren C. Holland, Jonathan E. Holme, T. Horch, P. Hrab, J. Hu, T.-H. Huynh, J.-Y. Hwang, R. Iacovelli, D. Iftime, M. Iorio, S. Jayachandran, E. Jeong, J. Jing, Jung J. Jung, Y. Kakumu, E. Kalkreuter, K. B. Kang, S. Kang, W. Kim, G. J. Kim, H. Kim, Hyun U. Kim, M. Klapper, Robert A. Koetsier, C. Kollten, Ákos T. Kovács, Y. Kriukova, N. Kubach, Aditya M. Kunjapur, Aleksandra K. Kushnareva, A. Kust, J. Lamber, M. Larralde, Niels J. Larsen, Adrien P. Launay, N.-T.-H. Le, S. Lebeer, B. T. Lee, K. Lee, Katherine L. Lev, S.-M. Li, Y.-X. Li, C. Licona-Cassani, A. Lien, J. Liu, Julius Adam V. Lopez, Nataliia V. Machushynets, Marla I. Macias, T. Mahmud, M. Maleckis, A. M. Martinez-Martinez, Y. Mast, Marina F. Maximo, Christina M. McBride, Rose M. McLellan, K. M. Bhatt, C. Melkonian, A. Merrild, M. Metsä-Ketelä, Douglas A. Mitchell, Alison V. Müller, G.-S. Nguyen, Hera T. Nguyen, Timo H J. Niedermeyer, Julia H. O'Hare, A. Ossowicki, Bohdan O. Ostash, H. Otani, L. Padva, S. Paliyal, X. Pan, M. Panghal, D. S. Parade, J. Park, J. Parra, M. P. Rubio, Huong T. Pham, Sacha J. Pidot, J. Piel, B. Pourmohsenin, M. Rakhmanov, S. Ramesh, Michelle H. Rasmussen, A. Rego, R. Reher, Andrew J. Rice, A. Rigolet, A. Romero-Otero, Luis R. Rosas-Becerra, Pablo Y. Rosiles, A. Rutz, B. Ryu, L.-A. Sahadeo, M. Saldanha, L. Salvi, E. Sánchez-Carvajal, C. Santos-Medellin, N. Sbaraini, Sydney M. Schoellhorn, C. Schumm, L. Sehnal, N. Selem, Anjali D. Shah, Tania K. Shishido, S. Sieber, V. Silviani, G. Singh, H. Singh, N. Sokolova, Eva C. Sonnenschein, M. Sosio, Sven T. Sowa, K. Steffen, E. Stegmann, Alena B. Streiff, A. Strüder, F. Surup, T. Svenningsen, D. Sweeney, J. Szenei, A. Tagirdzhanov, B. Tan, Matthew J. Tarnowski, Barbara R. Terlouw, T. Rey, Nicola U. Thome, L. R. Torres Ortega, T. Tørring, M. Trindade, Andrew W. Truman, M. Tvilum, Daniel W. Udwary, C. Ulbricht, L. Vader, Gilles P. van Wezel, M. Walmsley, R. Warnasinghe, Heiner G. Weddeling, Angus N. M. Weir, K. Williams, Sam E. Williams, Thomas E. Witte, Steffaney M. W. Rocca, K. Yamada, D. Yang, D. Yang, J. Yu, Z. Zhou, N. Ziemert, L. Zimmer, A. Zimmermann, C. Zimmermann, Justin J. J. van der Hooft, Roger G. Linington, T. Weber, Marnix H. Medema, *Nucleic*

*Acids Res* **2024**, *53* (D1), D678, https://doi.org/10.1093/nar/gkae1115; b) B. R. Terlouw, K. Blin, J. C. Navarro-Muñoz, N. E. Avalon, M. G. Chevrette, S. Egbert, S. Lee, D. Meijer, M. J. J. Recchia, Z. L. Reitz, J. A. van Santen, N. Selem-Mojica, T. Tørring, L. Zaroubi, M. Alanjary, G. Aleti, C. Aguilar, S. A. A. Al-Salihi, H. E. Augustijn, J. A. Avelar-Rivas, L. A. Avitia-Domínguez, F. Barona-Gómez, J. Bernaldo-Agüero, V. A. Bielinski, F. Biermann, T. J. Booth, V. J. Carrion Bravo, R. Castelo-Branco, F. O. Chagas, P. Cruz-Morales, C. Du, K. R. Duncan, A. Gavriilidou, D. Gayrard, K. Gutiérrez-García, K. Haslinger, E. J. N. Helfrich, J. J. J. van der Hooft, A. P. Jati, E. Kalkreuter, N. Kalyvas, K. B. Kang, S. Kautsar, W. Kim, A. M. Kunjapur, Y. X. Li, G. M. Lin, C. Loureiro, J. J. R. Louwen, N. L. L. Louwen, G. Lund, J. Parra, B. Philmus, B. Pourmohsenin, L. J. U. Pronk, A. Rego, D. A. B. Rex, S. Robinson, L. R. Rosas-Becerra, E. T. Roxborough, M. A. Schorn, D. J. Scobie, K. S. Singh, N. Sokolova, X. Tang, D. Udwary, A. Vigneshwari, K. Vind, S. Vromans, V. Waschulin, S. E. Williams, J. M. Winter, T. E. Witte, H. Xie, D. Yang, J. Yu, M. Zdouc, Z. Zhong, J. Collemare, R. G. Linington, T. Weber, M. H. Medema, *Nucleic Acids Res* **2023**, *51* (D1), D603, https://doi.org/10.1093/nar/gkac1049.
